# A conserved vesicular circuit enables metabolic adaptation and maintains systemic heme homeostasis

**DOI:** 10.64898/2026.07.31.742071

**Authors:** Sohini Dutt, Simon Beardsley, Xiaojing Yuan, Ran Yu, Audrey Belot, Blake Carrington, Kevin Bishop, Sandeepan Ghosh, Jiayu Li, Xuedi Zhang, Jean-Martin Harder, Rutwick Surya Ulhas, Shilpa Dilip Kumar, Vijaya Pandey, Raman Sood, Harold Smith, Michael Krause, James A Wohlschlegel, Ute A Hellmich, Caiyong Chen, Paul Liu, Iqbal Hamza

**Affiliations:** Center for Blood Oxygen Transport and Hemostasis, Department of Pediatrics, School of Medicine, University of Maryland, Baltimore, Maryland, USA; Department of Animal and Avian Sciences, University of Maryland, College Park, Maryland, USA; Zebrafish Core, National Human Genome Research Institute, National Institutes of Health, Bethesda, Maryland, USA; MOE Key Laboratory of Biosystems Homeostasis and Protection, College of Life Sciences, Zhejiang University, Hangzhou, China; Faculty of Chemistry and Earth Sciences, Institute of Organic Chemistry and Macromolecular Chemistry, Friedrich Schiller University Jena, Jena, Germany; Department of Pharmacology and Physiology, School of Medicine, University of Maryland, Baltimore, Maryland, USA; Department of Biological Chemistry, David Geffen School of Medicine, University of California Los Angeles, Los Angeles, California, USA; Laboratory of Molecular Biology, National Institute of Diabetes and Digestive and Kidney Diseases, National Institutes of Health, Bethesda, Maryland, USA; Cluster of Excellence, Balance of the Microverse, Friedrich Schiller University Jena, Jena, Germany; Oncogenesis and Development Section, Translational and Functional Genomics Branch, National Human Genome Research Institute, National Institutes of Health, Bethesda, Maryland, USA

## Abstract

Metabolic homeostasis depends on adaptive control of intracellular metabolite flux, yet how such control is reconfigured when canonical transport pathways fail is unknown. Here we define a conserved vesicular circuit that preserves systemic heme balance by rerouting intracellular heme flux. We show that loss of the intestinal heme exporter MRP-5 in *Caenorhabditis elegans* causes lethal heme sequestration within endolysosomal compartments. This defect is bypassed by disabling the vesicular adaptor AP-3, which stabilizes and reroutes the heme importer HRG-1, restoring heme export without increasing cytosolic heme. Unbiased genetics and transcriptomics identify two previously uncharacterized SLC49A family members, HRG-13 and HRG-14, as heme exporters with distinct affinities that engage in a vesicular importer-exporter handoff. Live imaging reveals heme-enhanced contacts between HRG-1 and SLC49A-containing vesicles, consistent with direct vesicular transfer. This circuitry extends to vertebrates as disruption of the SLC49A3 homolog impairs erythropoiesis in zebrafish and causes intracellular heme overload, premature hemoglobinization and apoptosis in differentiating human erythroid cells. Together, these findings establish SLC49A3 proteins as conserved heme exporters and uncover a general principle of metabolic adaptation in which reprogramming intracellular compartmentalization, rather than increasing nutrient supply, restores systemic homeostasis.

## Main

Organisms must adjust their metabolism in response to environmental changes and fluctuating nutrient availability through metabolic adaptations to maintain homeostasis ^1^. This adaptation depends on an organism’s ability to sense environmental shifts and adjust metabolic states. Genetic perturbations that disrupt essential nutrient pathways often cause severe growth defects that can be alleviated or bypassed by nutrient supplementation. However, the cellular mechanisms for this metabolite-based suppression, as described for iron, copper, zinc, and manganese ^2-9^, remain poorly understood.

Heme homeostasis provides a powerful system in which to dissect these mechanisms. *Caenorhabditis elegans*, like mammals, regulates its metabolism, growth, and behavior in response to heme. But unlike mammals, these free-living nematodes are unique as they lack the ability to synthesize heme and rely entirely on environmental heme rendering them sensitive to perturbations in heme acquisition and distribution ^10,11^. Heme, an iron-containing tetrapyrrole, serves as both a prosthetic group and a signaling molecule, regulating a wide range of biological processes ^12,13^. However, free heme is cytotoxic, as it integrates into lipid membranes and generates reactive oxygen species^14^. Consequently, heme trafficking between membrane compartments must be tightly regulated to balance supply with toxicity ^12^.

Dietary heme uptake in *C. elegans* is mediated by Heme Responsive Gene (HRG)-1-related transmembrane permeases ^15,16^. HRG-1 localizes to endolysosomal compartments in intestinal cells where it facilitates heme transport to the cytosol, whereas its paralog HRG-4 resides on the apical surface ^15,16^. Export of heme from the intestine to extra-intestinal tissues is mediated by the ABC transporter MRP-5/ABCC5 via the basolateral membranes ^17^. Loss of MRP-5 causes embryonic lethality due to a defect in systemic heme distribution. Importantly, this lethality can be suppressed by heme supplementation implying the existence of alternative heme export pathways ^17^.

To uncover these alternative pathways, we performed an unbiased genetic screen using the heme-dependent lethality of *mrp-5* mutants. We found that loss of subunits of the vesicular and cargo sorting complex Adaptor Protein-3 (AP-3) strongly suppressed *mrp-5* lethality. Disabling AP-3 rerouted HRG-1 dependent vesicular trafficking and restored intestinal heme efflux through HRG-13 and HRG-14, homologs of the previously uncharacterized vertebrate transmembrane protein SLC49A3. Our genetic, cellular, and biochemical studies in *C. elegans*, zebrafish and mammalian cells define a mechanistic model for metabolite-based suppression and reveal how regulated compartmentalization and vesicular trafficking allow organisms to overcome nutrient limitations while preserving systemic homeostasis.

## Results

### Metabolite-guided suppression identifies AP-3 inhibition bypasses MRP-5 driven heme export

We previously identified a role for MRP-5 in heme transport in both *C. elegans* and mammals ^17^. The *C. elegans mrp-5(ok2067)* mutants exhibit embryonic lethality, which can be rescued by providing excess dietary heme ^17^. However, recent studies proposed that increased intestinal membrane permeability and heme leakage, rather than a defect in heme export, gave rise to the heme-dependent phenotype of *mrp-5* mutants with high heme supplementation (≥ 500 µM) ^18^. To precisely establish the heme concentrations required to suppress the lethal phenotype, *mrp-5(ok2067)* mutants and their corresponding wildtype (WT) broodmates were cultured with increasing concentrations of heme in the growth medium (**Figure 1A**). Over 50% of embryos hatched and developed into larvae when exposed to ≥ 60 µM heme, indicating that the lethality in *mrp-5* mutants is due to heme deficiency. This result suggests that, in the absence of MRP-5, heme supplementation can exit the intestine through an alternative pathway.

**Figure 1.**
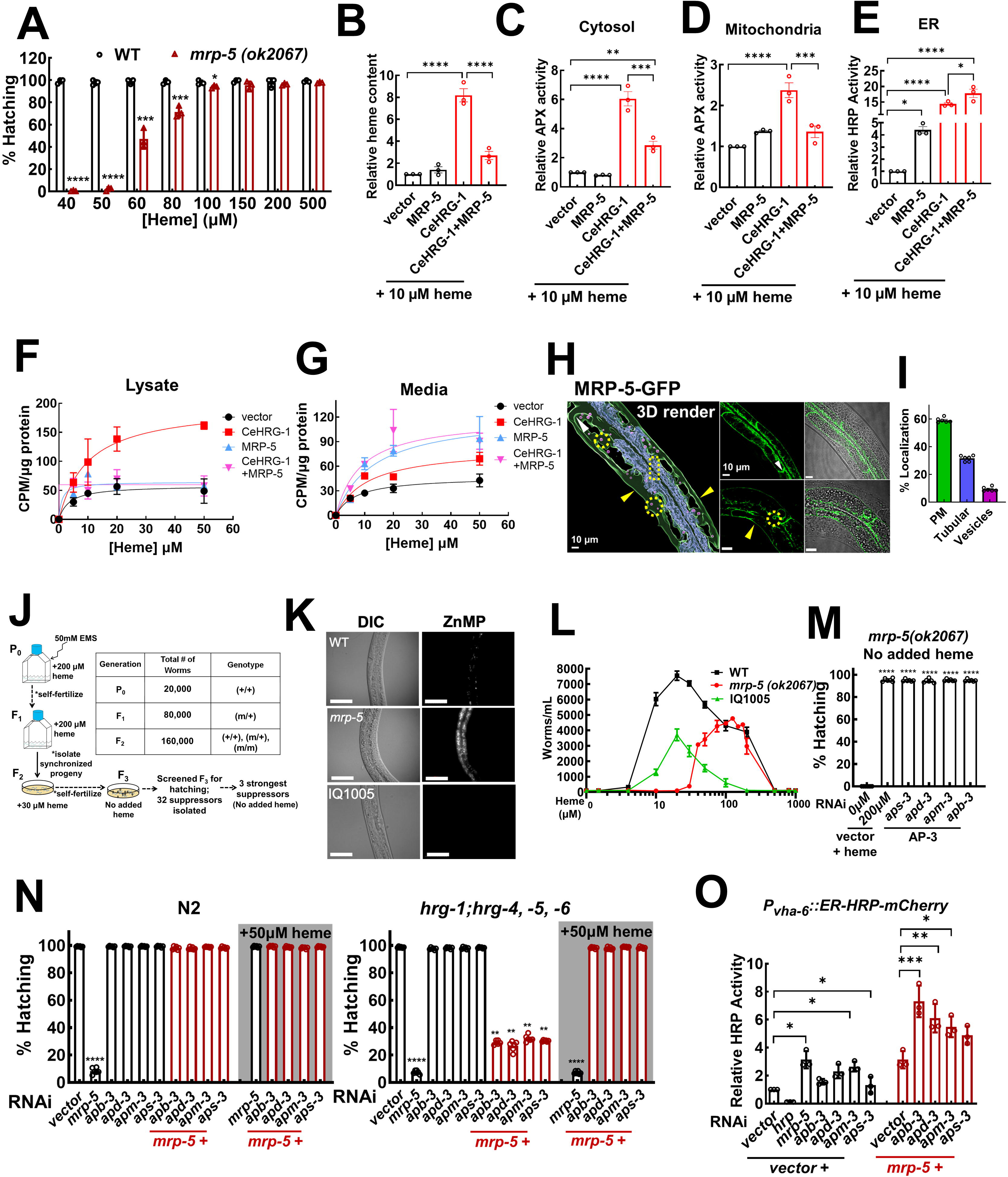
MRP-5 mediates heme export in *C. elegans*, and a forward genetic suppressor screen uncovered AP-3 complex subunits as strong suppressors of *mrp-5*-dependent lethality. (A) *mrp-5 (ok2067)* mutant worms show heme-dependent embryonic lethality (n=3). (B) Total heme content in HEK293 cells transiently transfected with the indicated constructs (n=3). Labile heme levels in the (C) cytosol, (D) mitochondria, and (E) endoplasmic reticulum (ER), were determined by expressing cytosol-APX, mitochondria-APX, and ER-HRP reporters, respectively (n=3). Transfected cells were supplemented with heme depleted (HD) serum + succinyl acetone (SA) + 10 µM heme. Error bars represent mean ± SEM from 3 biological replicates. [^55^Fe]heme measurements in (F) cell lysates and (G) media confirm that MRP-5 exports heme (n=2). (H) Representative confocal images of MRP-5 translation fusion worm strain shows that MRP-5 localization is distributed between the intestinal basolateral membrane (white arrowheads show lateral while yellow arrowheads show basal localization), tubular structures (shown by yellow boxes), as well as endosomal vesicles (shown by yellow circles). Left panel shows a 3D rendering of MRP-GFP in worm intestine, while the four right panels are representative confocal images. Scale bar,10 µm. (I) Quantification of MRP-5 localization across different cellular compartments: plasma membrane (PM), tubular, and vesicular compartments within an intestinal cell of six independent worms. (J) Schematic of the EMS mutagenesis screen conducted on *mrp-5(ok2067)* worms grown axenically in mCeHR-2 media supplemented with 200 µM heme. Following EMS treatment, mutagenized P_0_ animals were allowed to lay F_1_ progeny. Gravid F_1_ were bleached to isolate and synchronize the F_2_ generation, which were then seeded on plates supplemented with 30 µM heme. Only lines capable of laying viable progeny after seven days were called *mrp-5* suppressors. (K) *mrp-5* suppressor mutants alter ZnMP accumulation within the intestine. Indicated worm strains were grown and treated with ZnMP. Scale bar, 50 µm. (L) Indicated worm strains were bleached, and 100 synchronized L_1_ progeny were grown in mCeHR-2 media supplemented with increasing concentrations of heme for nine days and quantified (worms/ml) by microscopy (n=3). (M) RNAi depletion of AP-3 subunits rescues embryonic lethality caused by loss of *mrp-5* without the addition of any exogenous heme. *mrp-5(ok2067)* seeded on 200 µM heme was used as a positive control (n=6). (N) Loss of AP-3 subunits act as strong suppressors of *mrp-5* lethality. WT (N2) and quadruple mutant (*hrg-1; hrg-4,-5,-6,* QKO) were fed with dsRNA against *mrp-5* or ap-3 subunits or both in the presence and absence of 50 µM added heme (n=6). (O) Peroxidase reporter activity analysis of synchronized L_1_s of *P_vha-6_::ER-HRP::mCherry* seeded on indicated RNAi plates, harvested at young adult stage, lysed, and then subjected for activity measurements using o-dianisidine and H_2_O_2_ as substrates (n=3). *p<0.05, **p<0.01, ***p<0.001 ****p<0.0001 (one-way ANOVA with Dunnett’s multiple comparison test). Data shown as mean ± SEM.

To investigate heme export at the cellular level, we utilized HEK293 mammalian cells expressing worm MRP-5, both with and without the *C. elegans* heme importer HRG-1 (CeHRG-1) **(Supplemental Data Figure 1A)**. HRG-1 expression significantly increased total heme accumulation by nearly eight-fold, which was markedly reduced when MRP-5 was co-expressed (**Figure 1B, Supplemental Data Figure 1B**). To analyze subcellular heme distribution, we employed genetically encoded, peroxidase-based heme reporters ^19^. Horseradish and ascorbate peroxidases, which require heme as a cofactor for their activity, were targeted to various subcellular organelles to interrogate the labile heme availability within that compartment. Reduced cytosolic and mitochondrial peroxidase activity was observed in cells co-expressing MRP-5 (**Figure 1C and D**, **Supplemental Data Figure 1C and 1D**), while increased activity was detected in the endoplasmic reticulum (ER)/secretory pathway (**Figure 1E and Supplemental Data Figure 1E**).

To directly confirm heme export activity, we employed radioactive [^55^Fe]heme tracer studies. Conditions were first optimized in HEK293 cells to ensure heme loading by expressing CeHRG-1 (**Supplemental Data Figure 1F**). A time-dependent sustained heme uptake was observed. To measure heme efflux, cells co-expressing CeHRG-1 and MRP-5 were first pulse-labeled with [^55^Fe]heme, followed by a chase. A significant decrease in intracellular [^55^Fe]heme and a corresponding increase in extracellular media was observed in cells expressing MRP-5 with an apparent K_m_ of ≈8.7 µM (**Figure 1F and G**). These findings are consistent with the cellular localization of MRP-5 to the trans-Golgi compartment, intracellular vesicles/tubules, and plasma membrane in both mammalian cells and *C. elegans* (**Figure 1H and I**, **Supplemental Data Figure 1G and H**). Collectively, these results provide strong evidence that MRP-5 plays a direct role in heme export and facilitates its loading into the secretory pathway.

Since heme supplementation suppresses *mrp-5* embryonic lethality, we conducted an EMS-based forward genetic screen to identify alternative heme exporters. We screened approximately 160,000 haploid genomes, achieving eight-fold genomic coverage, to uncover suppressors of *mrp-5(ok2067)* lethality (**Figure 1J**). This screen identified three strong suppressors that produced viable F_3_ progeny without requiring additional heme supplementation (**Supplemental Data Figure 1I**). *mrp-5* mutants not only accumulate zinc mesoporphyrin IX (ZnMP), a fluorescent heme analog, in the intestine but also demonstrate resistance to the toxic heme analog gallium protoporphyrin IX (GaPPIX) indicating that heme analogs are able to enter the intestine but not efficiently transported to extraintestinal cells (**Figure 1K and Supplemental Data Figure 1K**) ^11,15^. Interestingly, all three suppressors displayed little to no detectable ZnMP signal indicating altered heme transport and were sensitive to GaPPIX toxicity phenocopying WT (**Figure 1K and Supplemental Data Figure 1J and K**). In mCeHR-2 liquid axenic medium, WT worms thrive at 20 µM heme but experience growth inhibition at heme concentrations ≥800 µM (**Figure 1L**) ^11^. By comparison, *mrp-5(ok2067)* mutants required 125 µM heme for optimal growth, while the suppressor strain (*IQ1005*) displayed a growth optimum at 20 µM heme. Notably, the suppressor exhibited sensitivity to high heme levels, with growth inhibition observed at 200 µM heme. Pre-culturing parental worms (P_0_) at varying heme concentrations did not alter the heme-dependent growth profiles of the *mrp-5*-deficient strain, ruling out maternal heme loading as a contributing factor (**Supplemental Data Figure 1L**) ^20^. These findings suggest that intestinal heme efflux is genetically restored in the suppressors, compensating for the loss of MRP-5.

Variant-based deep sequencing of the three bypass suppressors identified ≈2 Mb intervals on three different chromosomes (**Supplemental Data Figure 1M**). Remarkably, among the candidate mutations in each suppressor strain, we identified distinct subunits (*aps-3*, *apd-3*, and *apm-3*) of a single complex, Adaptor Protein 3 (AP-3) ^21,22^. RNAi depletion of other candidate genes within the mapped intervals in *mrp-5(ok2067*) mutants or *mrp-5* (RNAi) confirmed that loss of AP-3 subunits specifically mimicked *mrp-5* suppressor phenotypes (**Supplemental Data Figure 1N-Q**). Underscoring the importance of the AP-3 complex, depletion of *apb-3*, the fourth subunit of AP-3, also suppressed *mrp-5* lethality **(Figure 1M**).

To assess the strength of *mrp-5* suppression by loss of AP-3, we developed a heme-sensitized strain by deleting all four paralogs of *hrg-1* heme importers (*hrg-1*, *hrg-4*, *hrg-5*, and *hrg-6*). In these quadruple knockout (QKO) worms, loss of *mrp-5* caused embryonic lethality that could not be rescued by heme supplementation unless AP-3 subunits were co-depleted (**Figure 1N**).

Given that MRP-5 and AP-3 are expressed across multiple tissues, it is plausible that their depletion in specific tissues contributes to the suppression phenotype. Using tissue-specific RNAi strains, we found that only intestine-specific RNAi *(VP303)* suppressed *mrp-5* lethality (**Supplemental Data Figure 1R**). To evaluate whether AP-3 suppression was specific to heme, we assessed AP-3 deficiency in copper metabolism. CUA-1 is the *C. elegans* homolog of copper-exporting ATP7 that traffics from the secretory pathway to basolateral membranes to export intestinal copper; its loss is lethal but can be rescued partially by copper supplementation (**Supplemental Data Figure 1S**) ^23,24^. Co-depletion of *cua-1* and *apb-3* resulted in complete lethality, regardless of copper supplementation (**Supplemental Data Figure 1S**). Furthermore, exposing *mrp-5(ok2067)* worms to Dynasore and chloroquine, inhibitors of vesicle formation and endosomal acidification, rescued *mrp-5* associated lethality at 30 μM heme **(Supplemental Data Figure 1T)** ^25,26^.

We next investigated heme status in intestinal and extra-intestinal tissues, by employing genetically-encoded peroxidase reporters driven by tissue-specific promoters ^19^. ER-targeted HRP reporters revealed that loss of *mrp-5* significantly increased intestinal heme levels (**Figure 1O**), while reducing hypodermal and neuronal levels (**Supplemental Data Figure 1U,V**), indicating defective intestinal heme export. Strikingly, co-depletion of *mrp-5* and *AP-3* subunits elevated reporter activity across all three tissues ((**Figure 1O** and **Supplemental Data Figure 1U,V)**. Since ER-HRP reflects the heme status of the secretory pathway, we utilized a cytosol-targeted APX reporter driven from an intestinal promoter. RNAi of *mrp-5* with or without AP-3 subunits did not affect intestinal cytosolic heme levels (**Supplemental Data Figure 1W**). Collectively, our findings indicate that loss of AP-3 subunits in *mrp-5* mutants enhances heme within the intestinal secretory pathway accompanied by increased extra-intestinal heme levels.

### AP-3 loss alleviates heme-induced HRG-1 degradation in *mrp-5* mutants

AP-3 subunits play a role in sorting cargo proteins to lysosome-related organelles (LROs), and the heme importer HRG-1 localizes to the endolysosomes ^15,21,22^. To investigate whether AP-3 influences HRG-1 localization, we performed confocal microscopy on *hrg-1-gfp* transgenic worms fed with the fluorescent heme analog ZnMP ^15,17^. In WT worms, HRG-1 colocalized with both ZnMP and LysoTracker **(Figure 2A)**. RNAi depletion of *mrp-5* led to ZnMP accumulation in LysoTracker-positive compartments but, strikingly, the worms showed little to no visible HRG-1 signal (**Figure 2A**). Loss of *apb-3* resulted in reduced LysoTracker and ZnMP staining, along with diffused HRG-1 localization. Notably, these defects were effectively rescued by co-depleting *apb-3* and *mrp-5* (**Figure 2A**). Yeast three-hybrid (Y3H) assay using CeHRG-1 as bait confirmed a direct interaction between HRG-1 and AP-3 components, thus strengthening the genetic and localization relationship between HRG-1 and AP-3 (**Figure 2B**). To determine whether the loss of HRG-1 in *mrp-5* mutants is linked to heme accumulation, we analyzed two strains: one expressing HRG-1 from its endogenous promoter (*P_hrg-1_::hrg-1-gfp*) and another expressing HRG-1 ectopically from a strong intestinal promoter (*P_vha-6_::hrg-1-HA::sl2::mCherry*). In both cases, increased heme levels inversely correlated with HRG-1 abundance suggesting that heme accumulation is detrimental to HRG-1 protein (**Figure 2C**).

**Figure 2.**
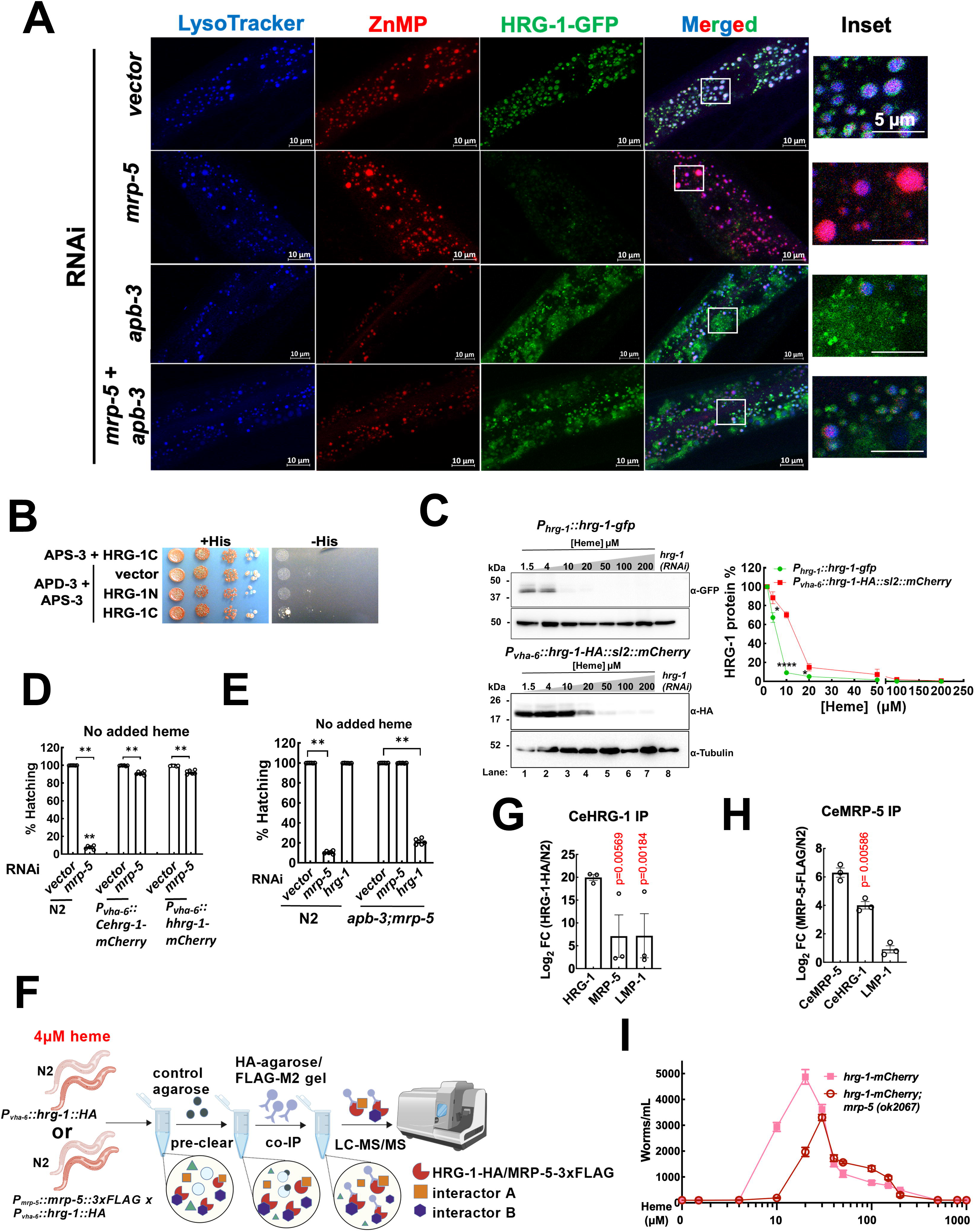
The heme importer HRG-1 dictates *mrp-5* lethality. (A) Coordinated regulation of *mrp-5* and AP-3 dictates HRG-1 abundance. Representative confocal microscope images showing *P_hrg-1_::HRG1-1::GFP* fed with indicated dsRNA, co-stained with LysoTracker Blue and Zinc mesoporphyrin (ZnMP). Scale bars as indicated. White boxes indicate insets, scale bar 5 µm. (B) Yeast three-hybrid (APD-3 fused with NLS and DNA activation domain, APS-3 fused with NLS, HRG-1 N-/C-terminus fused with NLS and DNA binding domain, to test interaction between APD-3/APS-3 hemicomplex and HRG-1) assay showed that CeHRG-1 interacts with AP-3 complex (C) Immunoblot analysis reveal heme-dependent turnover of HRG-1 protein. *P_hrg-1_::hrg-1-gfp* and *P_vha-6_::hrg-1-HA::sl2::mCherry* worms were grown axenically under different heme concentrations. Blots from three independent experiments were quantified using ImageJ (n=3). (D) HRG-1 levels dictate *mrp-5* lethality. *Cehrg-1* denotes *C. elegans* HRG-1, while *hhrg-1* represents worm expressing human HRG1 (n=6). (E) RNAi knockdown of *hrg-1* in suppressor strain *apb-3;mrp-5* reverses the suppression, resulting in lethality unlike WT worms (n=6). (F) Schematic showing co-immunoprecipitation (co-IP) workflow, made using BioGDP. IP of worm (G) HRG-1and (H) MRP-5 showing enrichment of indicated proteins. The p values here indicate tryptic peptides that were enriched after mass spectrometry analysis of co-IP eluates. (I) Heme-dependent growth curves of worms expressing CeHRG-1-mCherry (*vha-6p::Cehrg-1-mCherry*) in both WT and *mrp-5* KO background. Overexpressing CeHRG-1-mCherry did not change optimal heme requirement (20µM) in WT background. In *mrp-5(ok2067)* background, overexpressing CeHRG-1-mCherry restored optimal heme requirement from 125µM (by *mrp-5* KO) to 30µM, indicating additional heme transporter(s) are involved in rescuing the suppressor strains (to 20µM) (n=3). Error bars represent mean ± SEM. *p<0.05, **p<0.01, ***p<0.001 (one-way ANOVA with Dunnett’s multiple comparison test).

To directly assess whether *hrg-1* functions as a genetic modifier of *mrp-5*, we used transgenic worms expressing *C. elegans* (*Cehrg-1*) or human HRG-1 (*hhrg-1*) under the control of an intestine-specific promoter (*P_vha-6_::hrg-1::mCherry*). RNAi depletion of *mrp-5* in these transgenic worms did not cause lethality (**Figure 2D**). Conversely, depletion of *hrg-1* in the suppressor strain (*apb-3;mrp-5*) caused embryonic lethality, confirming a critical role for HRG-1 in the suppressor mechanism (**Figure 2E**). To further test whether HRG-1 and MRP-5 physically interact, we performed two complementary proteomic pull-downs (**Figure 2F**). Co-immunoprecipitation (IP) of HRG-1-HA followed by mass-spectrometry identified MRP-5 as one of its most enriched interactors (**Figure 2G**). Reciprocally, co-IP of worms expressing MRP-5-FLAG recovered HRG-1-HA as one of the top interactors of MRP-5 (**Figure 2H**). Consistent with this, RNAi knockdown of *ap-2* and *ap-3* in *hrg-1* and *mrp-5* reporter strains caused mis-localization of both proteins (**Supplemental Data Table 2**). Overexpression of HRG-1 in the *mrp-5* null background not only rescued overall worm growth, but shifted the optimal heme concentration for viability from 125 µM to 30μM supporting a dose-dependent role of HRG-1 in suppressing *mrp-5* lethality (**Figure 2I)**. Together, these findings demonstrate that loss of MRP-5 causes intestinal heme accumulation and consequently HRG-1 degradation.

### AP-3 loss reveals SLC49A transporters as alternative heme exporters

To elucidate the mechanisms by which AP-3 loss restores heme export in the absence of MRP-5, we performed differential gene expression analyses using RNA-seq data from WT, *mrp-5*, *apb-3*, and *apb-3;mrp-5* worms. A strong negative correlation was observed between differentially expressed genes (DEGs) in *mrp-5* and *apb-3;mrp-5* mutants, while no correlation was evident between *apb-3* and *apb-3;mrp-5* mutants (**Figure 3A and B**). Heatmaps revealed that previously characterized HRGs were significantly upregulated in *mrp-5* mutants compared to WT worms, but this upregulation was notably reduced in the suppressor strain (**Supplemental Data Figure 2A**) ^27^. Among the 3308 DEGs upregulated in the suppressor strain, 220 were predicted to encode proteins with more than one transmembrane domain (**Supplemental Data Figure 2B**). Gene set enrichment analysis identified consistent activation of genes involved in metabolite transport **(Figure 3C).** One of the most significant DEGs belonged to the *SLC49A* family (**Supplemental Data Figure 2C-E**).

**Figure 3.**
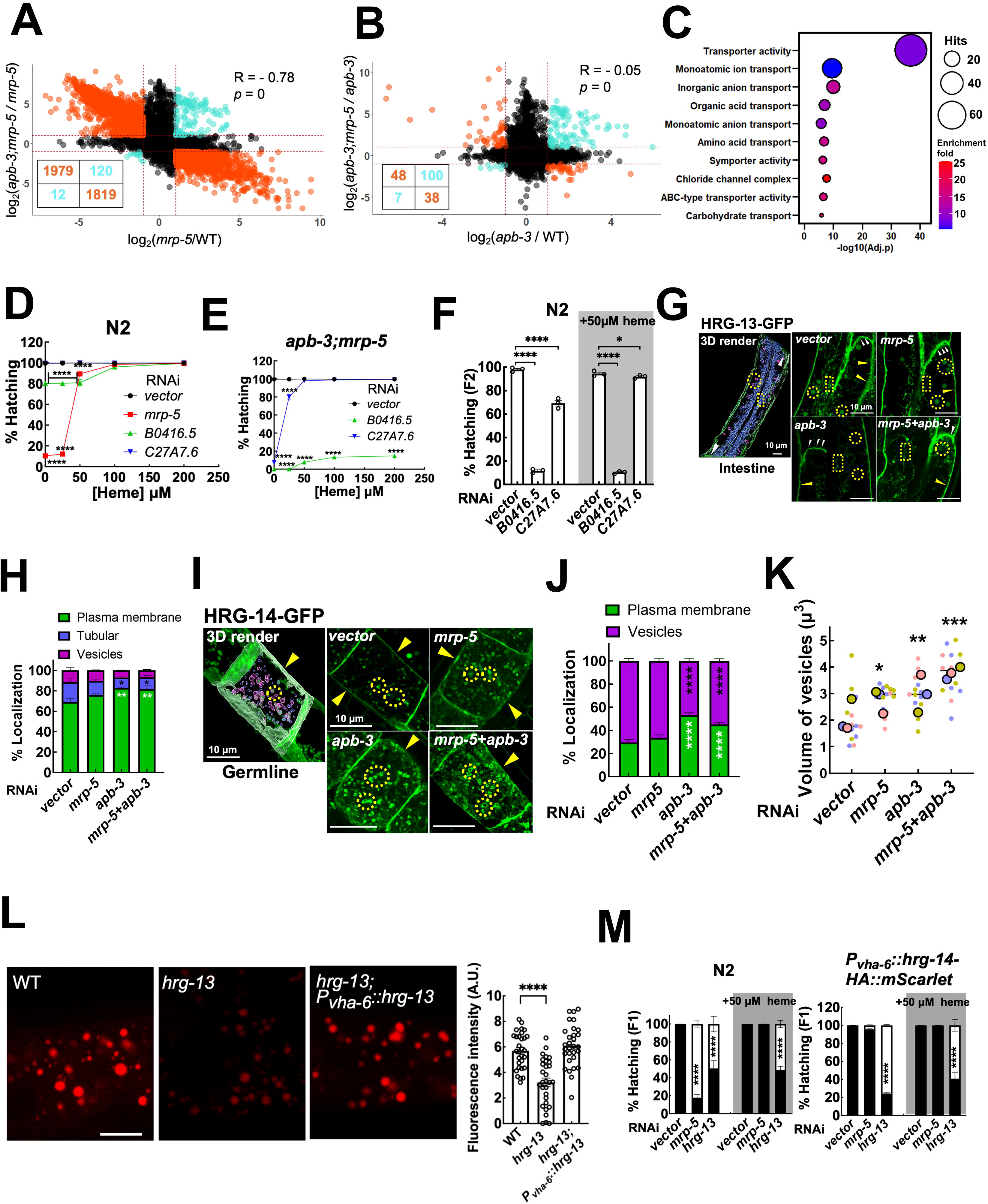
RNA-Seq identifies *SLC49A* homologs as alternative heme transporters. Scatter plots show differentially expressed transcripts as revealed by RNA-Seq analysis. (A) Plot log_2_ fold change (LFC) of *mrp-5(ok2067)* vs. WT (x-axis) against LFC of *mrp-5(ok2067);apb-3(ok429)* vs. *mrp-5(ok2067)* (y-axis). (B) Plot LFC of *apb-3(ok429)* vs. WT (x-axis) against LFC of *mrp-5(ok2067);apb-3(ok429)* vs. *apb-3(ok429)* (y-axis). Upper right and lower left quadrants contain positively co-regulated transcripts (cyan). Upper left and lower right quadrants contain negatively co-regulated transcripts (orange). Co-regulated transcripts had an absolute LFC change higher than 1 in both pairwise comparisons. Broken red lines marks LFC = 1. Correlation analysis was performed using Pearson method (Top: R=-0.78, Bottom: R=-0.05). RNA-Seq was performed using three biological replicates. (C) Gene Ontology (GO) analysis plots of the 220 genes that have more than one TMD. Heme dose response assays after RNAi depleting *B0416.5* and *C27A7.6* in (D) N2 and (E) *apb-3;mrp-5* (n=3). (F) RNAi knockdown of *hrg-13* and *hrg-14* show an enhanced phenotype in the F_2_ generation in N2 worms (n=3). (G) Left panel shows a 3D rendered image of HRG-14-GFP (*vector* RNAi) within the worm intestine. Representative confocal images showing HRG-14 localization in translational fusion worm strain subjected to RNAi knockdown as indicated on top left for each condition. Yellow arrowheads show basal surface, white arrowheads show lateral surface, yellow circles show intracellular vesicles, and yellow boxes show tubulations. (H) Quantification of HRG-13 localization across different cellular compartments within the intestine. White * and black * indicate statistical significance for plasma membrane and tubular compartments respectively. One intestinal cell from five independent worms were analyzed for quantification. (I) Left panel shows 3D rendered image of HRG-14-GFP (*vector* RNAi) within the worm germline. Representative confocal images showing HRG-14 localization in translational fusion worm strain subjected to RNAi knockdown as indicated on top left for each condition. Yellow arrowheads show plasma membrane localization, and yellow circles show intracellular vesicles. Within the 3D rendered image, vesicles less than 0.1 µm in diameter have been colored as blue, while those greater than 0.1 µm are shown in magenta. Scale bar, 10 µm. (J) Quantification of HRG-14 localization within different cellular compartments. White * and black * indicate statistical significance for plasma membrane and vesicular compartments respectively. At least four cells from three independent worms were quantified. (K) Quantification of volume of the vesicles within each cell in the germline. Each small circle represents one cell. Bigger circle of the same color represents mean of the volume of vesicles. At least four cells from three independent worms were quantified. (L) Representative images and quantification of ZnMP uptake assay in indicated worm strains. Scale bar, 10 µm. (M) Lethality assay in indicated synchronized worm strains in the presence and absence of added heme, scored for number of hatched worms and unhatched embryos. Error bars represent mean ± SEM. *p<0.05, **p<0.01, ***p<0.001, ****p<0.0001 (one-way ANOVA with Dunnett’s multiple comparison test).

The *SLC49A* family, part of the major facilitator superfamily, includes four human paralogs (**Supplemental Data Figure 2C**). While previous research established roles for human *SLC49A1* (FLVCR1) and *SLC49A2* (FLVCR2) in heme transport ^28-32^, recent studies implicate them to import choline and ethanolamine ^33-37^. Functional studies for the other two human paralogs, *SLC49A3* and *SLC49A4*, are lacking ^38^. The *C. elegans* genome encodes eight potential SLC49A homologs. Six were abundantly expressed in WT worms, whereas mRNA levels for *Y43D4A.3* and *Y43D4A.4* were barely detectable (**Supplemental Data Figure 2D and E**). In silico predictions of the six homologs indicated that all encoded full-length SLC49A with 12 transmembrane domains ^38^, while Y43D4A.3 and Y43D4A.4 appeared to be truncated versions of C27A7.6.

To test whether *SLC49A* homologs contributed to the rescue of *mrp-5* lethality, we RNAi depleted each of the six full-length worm homologs. In the *apb-3;mrp-5* suppressor, depletion of *B0416.5* and *C27A7.6* caused near-complete lethality (**Supplemental Data Figure 2F and G**), while heme supplementation fully rescued the lethality caused by *C27A7.6* depletion but had minimal effect on *B0416.5* depletion (**Figure 3D and E**). Similarly, depleting *B0416.5* in *mrp-5* mutants resulted in complete lethality, which was partially rescued (< 50%) by 200 µM heme (**Supplemental Data Figure 2H**). Knockdown of either *B0416.5* or *C27A7.6* did not exacerbate phenotypes in the QKO strain (**Supplemental Data Figure 2I**). In WT worms, only *B0416.5* depletion caused modest but significant lethality (∼20%) in the F_1_ generation that was rescued by heme supplementation **(Supplemental Data Figure 2F and Figure 3D, F**). However, this lethality was more pronounced in the F_2_ generation, with embryonic lethality observed for both *B0416.5* (>80%) and *C27A7.6* (>30%) (**Figure 3F**). Given that human *SLC49A1* and *SLC49A2* has been implicated in choline import ^33-37^, we investigated whether choline supplementation could rescue the phenotypes caused by *SLC49A* depletion in WT, *apb-3;mrp-5*, *QKO*, or *mrp-5* strains. Choline supplementation had no effect on phenotypes caused by SLC49A knockdowns (**Supplemental Data Figure 2J-M)**. We termed *B0416.5* and *C27A7.6* as *hrg-13* and *hrg-14*, respectively.

To determine the temporospatial localization of HRG-13 and HRG-14, we used CRISPR/Cas9 to generate C-terminal GFP fusions at their endogenous loci. HRG-13-GFP was expressed ubiquitously across developmental stages and localized to both intracellular puncta and basolateral membranes in intestinal cells, resembling MRP-5 localization (**Supplemental Data Figure 2N** and **1H)**. HRG-14-GFP was primarily expressed in the germline (**Supplemental Data Figure 2O**). Both proteins exhibited similar intracellular localization patterns, targeting the plasma membrane and vesicular/tubular compartments (**Figure 3G and I**). Depletion of *apb-3* with or without *mrp-5* disrupted the localization of both transporters. HRG-13 showed greater enrichment on the basolateral membranes (**Figure 3G and H**), while HRG-14 exhibited changes in both, the size/volume of the vesicles and enrichment on the plasma membrane (**Figure 3I-K**). To define factors regulating HRG-13 and HRG-14, we RNAi-depleted 40 genes encoding known membrane-trafficking components ^39^ in HRG-13-GFP and HRG-14-GFP strains (**Supplemental Data Table 3**). Knockdown of clathrin heavy chain (*chc-1*), COPI subunits (*copb-2, copg-2*), or vacuolar ATPase components (*vha-10*) produced aberrant reporter abundance or mislocalization. These findings indicate that, despite HRG-14’s germline restriction, both proteins rely on shared vesicular and membrane-trafficking regulators.

*hrg-13* knockout worms showed reduced intestinal ZnMP accumulation, consistent with its endogenous localization, and intestinally driven *hrg-13* expression restored ZnMP levels (**Figure 3L**). In contrast, *hrg-14* mutants displayed no change in intestinal ZnMP fluorescence (**Supplemental Data Figure 2P**), likely reflecting the germline-restricted expression of *hrg-14*. Notably, ectopic intestinal expression of *hrg-14* (P_vha-6_::*hrg-14-HA::mScarlet)* rescued *mrp-5*-associated lethality but not that caused by *hrg-13* depletion (**Figure 3M**). Together, these results indicate that in the absence of MRP-5, HRG-13 and HRG-14 act as alternative, yet functionally distinct, heme transporters.

### SLC49A3 homologs regulate heme balance and vertebrate erythropoiesis

We next assessed the ability of the *C. elegans* transporters and their corresponding human homologs to interact with heme. Using AlphaFold3 structural predictions ^40^ for the four human SLC49A homologs, we identified a putative binding site within the transmembrane cleft of only human SLC49A3 **(Figure 4A)**. Atomistic molecular dynamics (MD) simulations supported stable association of heme with SLC49A3 and HRG-14 **(Figure 4A-C, Supplemental Data Figure 3A-D)**. In the resulting SLC49A3-heme complex, the protoporphyrin ring is sandwiched between two aromatic residues, while the propionate groups form hydrogen bonds with hydroxyl-bearing side chains **(Figure 4B, Supplemental Data Figure 3A)**. The methyl/vinyl face of heme is accommodated in a hydrophobic groove formed by surrounding residues. This binding architecture is conserved in HRG-14 and MD simulations confirm a similar binding mode and conserved physicochemical properties of the heme-binding pocket **(Figure 4C, Supplemental Data Figure. 3A**), supporting functional complementarity between the two transporters.

**Figure 4.**
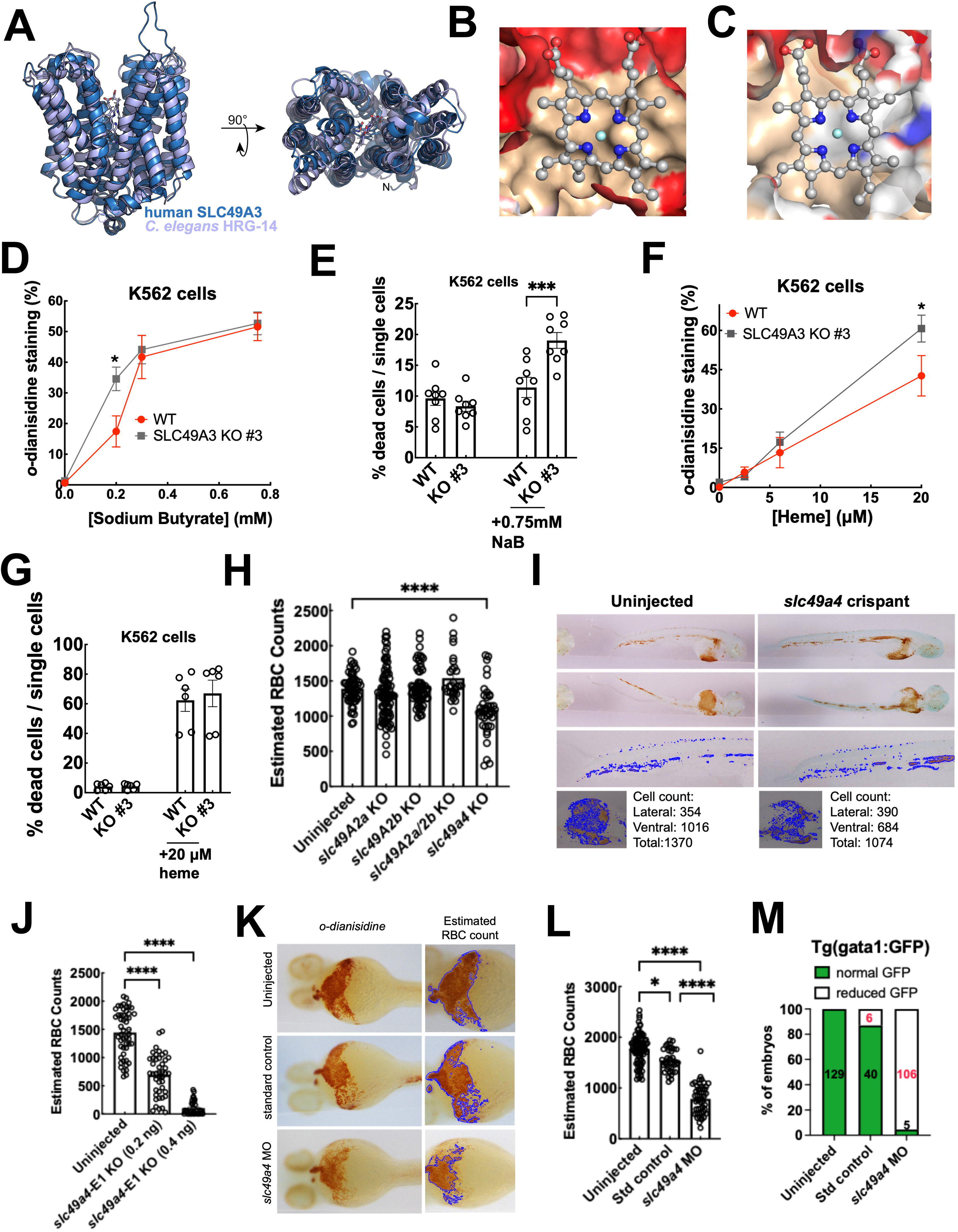
Conserved role of SLC49A family transporters in regulating heme balance and vertebrate erythropoiesis. (A) Overlay of human SLC49A3 (blue) and CeHRG-14 (light purple) structural snapshots from atomistic Molecular Dynamics Simulations carried out in an asymmetric eukaryotic plasma membrane (composition in **Supplemental Data Table 4**). For comparison of the conserved transmembrane core, the alignment of residues 25-443 of SLC49A3 and 43-468 of HRG-14 are shown yielding an all atom RMSD of 2.9 Å. For better visualization of the protein, the membrane, water and ions have been removed. (B) Surface representation of heme binding sites in SLC49A3 and (C) CeHRG-14 showing a similar distribution of hydrophobic (sand) residues contacting the protopophyrin ring and a charged region encasing the heme propionate groups. (D) WT and SLC49A3 KO cells were treated twice with different concentrations of sodium butyrate (NaB) and harvested 72h after second treatment. Cells were stained with benzidine, as an indirect method to quantify hemoglobinization. (E) Percentage of dead cells in WT and SLC49A3 KO cells with and without NaB treatment, when stained with viability dyes Annexin V and 7AAD. (F) WT and SLC49A3-KO cells were treated twice with different concentrations of hemin and harvested 72h after second treatment. Cells were stained with benzidine, as an indirect method to quantify hemoglobinization. (G) Percentage of dead cells in WT and SLC49A3 KO cells with and without NaB treatment, when stained with viability dyes Annexin V and 7AAD. (H) Quantification of total estimated RBC counts by treatment group. Embryos targeted with *slc49a4* showed significant reduction compared to uninjected controls. Sample sizes: Uninjected (n=55), *slc49A2a (*n=86), *slc49A2b* (n=62), *slc49A2a/2b* co-injection (n=28), and *slc49a4* (n*=*39). (I) Representative lateral and ventral images of 48 hpf zebrafish embryos injected with crispants and stained with *o-*dianisidine to visualize RBCs. Corresponding images analyzed using ImagePro software show gated regions (blue overlay) used for RBC quantification. (J) Dose-dependent reduction in RBC counts in embryos injected with *slc49a4* sgRNA targeting Exon 1. Sample sizes: uninjected (n=51), *slc49a4-*0.2ng (n=44), *slc49a4-* 0.4ng (n=35). (K) Representative images of 48 hpf embryos injected with 4800pg of *slc49a4* morpholino, stained with *o*-dianisidine, and analyzed by ImagePro software. Left: RBC staining. Right: gated overlay. (L) RBC quantification after morpholino injection. A slight reduction was observed in the standard control, whereas *slc49a4* morphants showed a strong reduction. Sample sizes: uninjected (n = 84), standard control 4800pg (n = 84), *slc49a4* MO 4800pg (n = 50). (M) GFP expression at 72 hpf in *Tg(gata1:GFP)* zebrafish injected with 4800pg of standard control or *slc49a4* morpholino. GFP levels were reduced in *slc49a4* morphants compared with controls.. Error bars represent mean ± SEM. *p<0.05, **p<0.01, ***p<0.001, ****p<0.0001 (one-way ANOVA with Dunnett’s multiple comparison test).

To assess the role of SLC49A3 in heme homeostasis, we used K562 human erythroleukemia cells, which differentiate toward the erythroid lineage upon sodium butyrate (NaB) treatment ^41-43^. Using CRISPR/Cas9, we generated multiple SLC49A3 knockout lines by deleting the region between exons 2 and 9 (KO#1, #3, #4, #5) and confirmed the precise lesions by sequencing each clone (**Supplemental Data Fig. 3E**). NaB-treated K562 cells accumulated increased heme and generated a higher proportion of benzidine-positive, hemoglobinized cells **(Figure 4D)**. In contrast, *SLC49A3*-deficient cells exhibited precocious differentiation, marked by significantly elevated heme accumulation compared to controls **(Figure 4D** and **Supplemental Data Figure 3F)**. Since NaB treatment induced premature hemoglobinization, we assessed apoptosis and observed higher 7AAD/AnnexinV-staining in NaB treated *SLC49A3-KO* cells (**Figure 4E** and **Supplemental Data Figure 3G**). Hemin treatment, another differentiation factor, produced similar effects with enhanced benzidine staining and elevated apoptosis in KO cells (**Figure 4F**, **G, Supplemental Data Figure 3H**, **I**).

To investigate the role of SLC49A in erythropoiesis, we used zebrafish as a vertebrate model system. Owing to the teleost-specific whole-genome duplication, direct one-to-one gene orthology is uncommon ^44^ **(Supplemental Data Figure 3J)**. However, AlphaFold structural modeling combined with *in silico* heme-binding analyses of the five zebrafish SLC49A homologs predicted heme binding within the transmembrane cleft of SLC49A2A, SLC49A2B, and SLC49A4 (DIRC2) **(Supplemental Data Figure 3K-M)**. CRISPR/Cas9-mediated disruption of all three paralogs in the zebrafish larvae (crispant) revealed only *slc49a4/dirc2* crispants with markedly reduced red blood cells (RBCs), as evidenced by diminished o-dianisidine staining (**Figure 4H, I**). A repeat of *slc49a4* crispants using the guide RNA targeting exon 1 at higher doses, elicited an even stronger reduction in RBCs. (**Figure 4J**). To independently validate these results, we performed morpholino (MO) knockdown of *slc49a4*. MO-treated embryos exhibited a robust, dose-dependent loss of o-dianisidine staining and reduced RBC numbers (**Figure 4K, L; Supplemental Data Figure 3N, O**). Correspondingly, MO injections into *Tg(gata1:GFP)* embryos, which expresses GFP in erythroid cells, markedly decreased *gata1* erythroid cells confirming that loss of *slc49a4* compromises erythropoiesis in vivo **(Figure 4M; Supplemental Data video 1)**. Collectively, these findings reveal a conserved role for SLC49A3 homologs in regulating erythropoiesis in vertebrates.

### HRG-13/14 drive cellular heme efflux through coordinated vesicular dynamics

To investigate whether HRG-13 and HRG-14 transport heme, we tagged these proteins with HA and GFP at their C-termini and ectopically expressed them in HEK293 cells (**Figure 5A**). Confocal microscopy confirmed that their localization in mammalian cells mirrored the patterns observed in *C. elegans*, including the plasma membrane and LAMP1-positive intracellular compartments. Notably, both proteins co-localized with CeHRG-1, suggesting a functional association between heme importers and exporters (**Figure 5B and C**).

**Figure 5.**
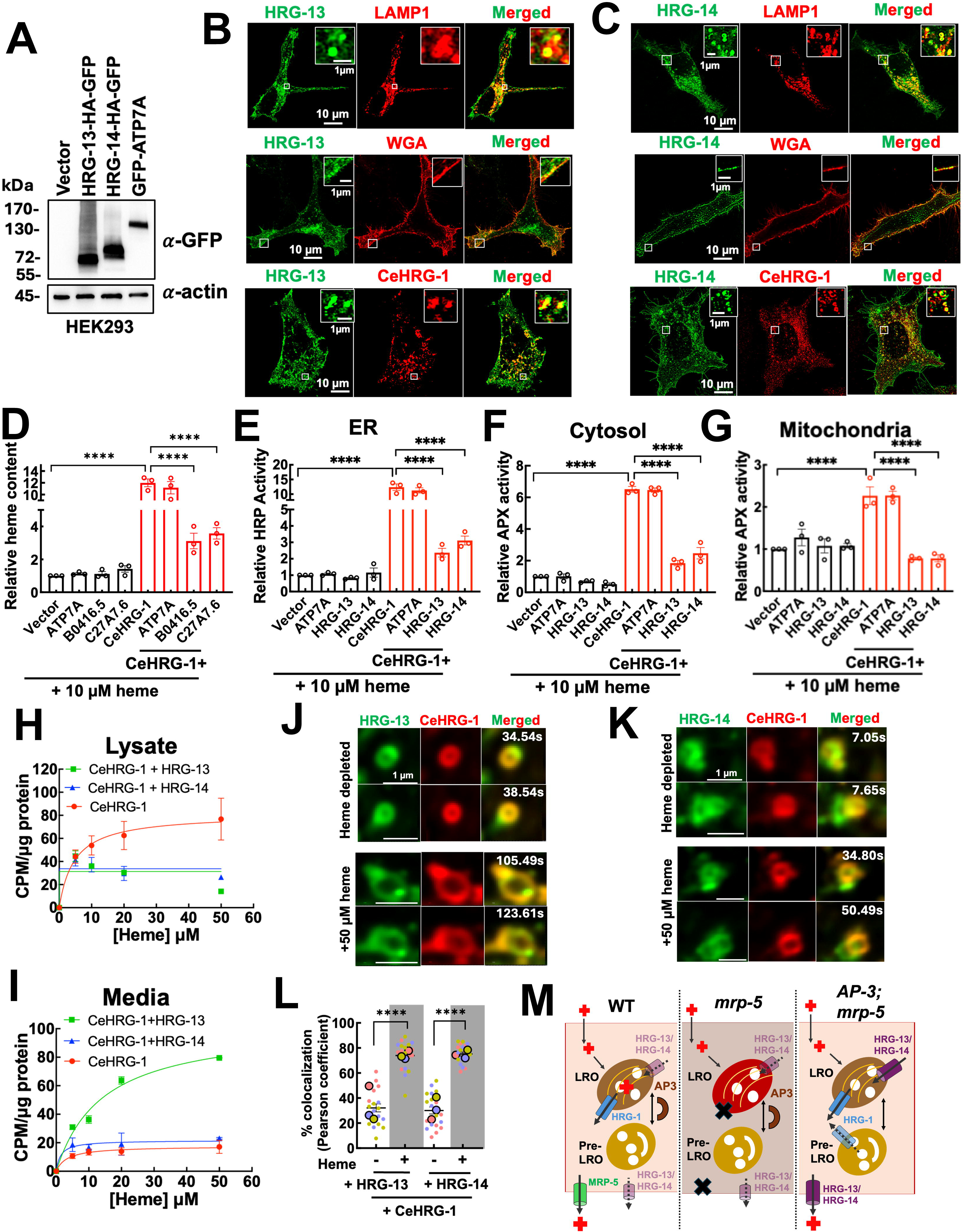
HRG-13 and HRG-14 export heme through vesicular coalescence with HRG-1. (A) Immunoblot analysis of HEK293 cells transiently transfected with indicated plasmids, probed with anti-GFP antibody. Representative confocal images showing HeLa cells transiently expressing (B) GFP-tagged HRG-13 and (C) HRG-14, along with LAMP-1, WGA, and CeHRG-1. White boxes represent insets. Scale bars as indicated. (D) Total heme content measurements show HRG-13 and HRG-14 reduce overall heme content (n=3). Subcellular labile heme measurements using peroxidase reporters show reduction in heme levels within the (E) ER, (F) cytosol, and (G) mitochondria (n=3). All heme content analysis were done under HD+SA+10 µM heme conditions. [^55^Fe]heme measurements in (H) cell lysates and (I) media confirm that HRG-13 and HRG-14 indeed function as heme exporters (n=2). Live-cell imaging using spinning-disk confocal microscope reveals vesicular coalescence between import and export vesicles under heme depleted and added conditions for both (J) HRG-13 and (K) HRG-14. (L) Quantification of % colocalization for cells expressing HRG-13 and HRG-14 with CeHRG-1 in the presence and absence of heme. Larger circles represent mean of all the replicates in the same color (smaller circles). At least five independent cells (represented in the same color) from three independent experiments were quantified for colocalization analysis. Pearson colocalization index was calculated across 3mins videos for each cell. Error bars represent mean ± SEM. *p<0.05, **p<0.01, ***p<0.001, ****p<0.0001 (one-way ANOVA with Dunnett’s multiple comparison test). (M) Proposed model for interplay between heme import, storage, and export.

To evaluate their role in heme export, we co-expressed CeHRG-1, which ensured that the cells are heme-loaded, with either HRG-13 or HRG-14. Cells co-transfected with CeHRG-1 and a heme exporter (HRG-13 or HRG-14) exhibited a significant reduction in total heme content compared to cells expressing CeHRG-1 alone (**Figure 5D, Supplemental Data Figure 4A**). Subcellular labile heme measurements using peroxidase-based reporters further demonstrated a decrease in reporter activity in all three major compartments (ER, cytosol, and mitochondria) of cells co-expressing either HRG-13 or HRG-14 with CeHRG-1 (**Figure 5E-G** and **Supplemental Data Figure 4B - D**). In contrast, control cells expressing the human copper exporter ATP7A showed no change in heme levels, confirming the specificity of HRG-13 and HRG-14 for heme export. We next assessed heme export activity in HEK293 cells co-expressing CeHRG-1 along with either HRG-13 or HRG-14, using [55Fe]heme as a tracer. A decrease in intracellular [^55^Fe]heme was observed alongside a corresponding increase in extracellular levels in cells expressing HRG-13 or HRG-14 (**Figure 5H and I, Supplemental Data 4F, G**). The apparent K_m_ values were ≈14.2 µM for HRG-13 and ≈1.4 µM for HRG-14. These results collectively indicate that both HRG-13 and HRG-14 enable heme efflux, reducing total cellular heme levels.

Live-cell imaging using spinning-disk confocal microscopy revealed dynamic interactions between vesicles containing CeHRG-1 and HRG-13 or HRG-14 (**Figure 5J and K**). Under low heme conditions, HRG-13 and HRG-14-containing vesicles transiently interacted with CeHRG-1 for < 4 s. But these interactions were significantly enhanced to > 15 s in the presence of heme suggesting vesicular coalescence as a mechanism for heme transfer between import and export compartments (**Figure 5L, Supplemental Data videos 2 to 5**). In summary, our findings demonstrate that HRG-13 and HRG-14 function as heme exporters, and that their co-localization with CeHRG-1 suggests a coordinated mechanism for heme transfer between heme importers and exporters by vesicular dynamics.

## Discussion

We identify SLC49A3 as a conserved metazoan heme exporter and place it within a dedicated importer-exporter circuit that organisms deploy to overcome genetically-imposed nutrient limitations. In *C. elegans*, loss of MRP 5 traps heme within LROs, triggers degradation of the endolysosomal importer HRG 1, and causes embryonic lethality (**Figure 5M**). This lethal bottleneck is relieved not only by dietary heme but, strikingly, by disabling the vesicular coat adaptor AP 3 ^21,22,45^, an intestine autonomous and heme specific suppression inconsistent with membrane leakage but consistent with regulated re routing. Extending this circuitry to vertebrates, we show that perturbing *slc49a4* compromises erythropoiesis in zebrafish, and loss of *SLC49A3* in differentiating human K562 cells elevates intracellular heme, accelerates hemoglobinization resulting in apoptosis. Together, these data elevate SLC49A3 from an uncharacterized solute carrier to a *bona fide* heme exporter that tunes cellular heme loads during development.

AP 3 acts through HRG 1 to set the balance between storage and export. HRG 1 contains tyrosine and di leucine based sorting motifs recognized by adaptor complexes such as AP 3 ^16,45^. When MRP 5 is absent, heme overload in LROs drives HRG 1 loss. Removing AP 3 stabilizes and missorts HRG 1, restores its localization, and re establishes heme flow through the secretory/basolateral pathway without elevating cytosolic heme, as shown by ER targeted sensors. Pharmacologic disruption of vesicle formation and endolysosomal acidification phenocopies the genetic rescue. The bypass is selective supporting a dedicated heme trafficking program rather than global membrane perturbation.

Unbiased suppression mapping and transcriptomics uncovered two *C. elegans* SLC49A family members, HRG 13 and HRG 14, as exporters that are recruited when AP 3 is lost. Endogenous tagging places HRG 13 at intestinal basolateral membranes and HRG 14 primarily in the germline; both depend on shared vesicular regulators and lie downstream of AP 3 by epistasis. In human cells heme loaded via HRG 1, HRG 13 and HRG 14 lower total and compartmental labile heme and drive heme efflux. Live cell imaging reveals heme enhanced, prolonged contacts between HRG 1 positive vesicles and HRG 13/14 compartments, suggesting a dynamic heme handoff at the importer exporter vesicular interface. Choline supplementation does not modify the in vivo phenotypes, separating these transporters from reported choline/ethanolamine activities of FLVCR1 and 2 ^33-37^.

Consistent with a direct export role, comparative modeling and in silico heme binding identify SLC49A3 as unique among human SLC49A paralogs, with a predicted cavity that accommodates heme between transmembrane helices, an architecture not evident for SLC49A1 and 2 (FLVCR1 and 2) ^28-30,38^,^31,32^. Together, the worm, zebrafish and human data identify SLC49A3 proteins as metazoan heme exporters that buffer heme to align supply with cellular and organismal demand. We propose that LRO vesicles serve as adaptive hubs where coat adaptors, luminal pH, and cargo specific transporters coordinate uptake, storage, and export ^46,47^, ^48,49^. Defining the steps that route HRG 1 for heme dependent turnover, the factors that partition HRG 13 and 14 across tissues, and the transport mechanism and stoichiometry of SLC49A3 will clarify how this circuit can be targeted to correct heme maldistribution in hematologic and mitochondrial disease. Our metabolite based suppression findings exemplify a general principle of nutrient driven plasticity: correcting compartmentalization, rather than increasing supply, can restore systemic homeostasis.

## Acknowledgments

We thank Juan Bonifacino, Xiaofei Bai, Andy Golden, and Michael Petris for reagents. We thank the Confocal Microscopy Core at the Center for Innovative Biomedical Resources, University of Maryland School of Medicine Baltimore; the use of the Zeiss LSM980 Aryscan 2 (supported by NIH 1S10OD025223) at the Imaging Core Facility at the University of Maryland College Park; HPC-cluster Draco at Universitätsrechenzentrum of Friedrich-Schiller-University Jena, Germany, and the HPC cluster of TU Ilmenau, Germany; and the *Caenorhabditis* Genetics Center for providing worm strains. The [^55^Fe] isotope used in this research was supplied by the U.S. Department of Energy Isotope Program, managed by the Office of Isotope R&D and Production. This work was supported by funding the National Institutes of Health DK125740 and DK074797 (to I.H.); German Research Foundation (Deutsche Forschungsgemeinschaft) EXC 2051 project ID 390713860 (to UAH); Cluster of Excellence Balance of the Microverse travel fellowship (to X.Z.); Intramural Research Program of the National Human Genome Research Institute, NIH to BK, KB, RS, PL. Their contributions are considered as works of the United States Government. Their findings and conclusions presented in this paper are those of the authors and do not necessarily reflect the views of the NIH or the US Department of Health and Human Services.

## Contributions

S.D., S.B., X.Y., P.L., C.C., and I.H. conceived the study, designed experiments, interpreted data, and wrote the manuscript. S.D., S.B., X.Y., R.Y., A.B., B.C., K.B., S.G., J.L., X.Z., V.P., performed experiments. S.D. and S.D.K. performed image analyses and quantifications. J.M.H., R.S.U., and U.A.H. performed and analyzed Molecular Dynamics Simulations and protein structure predictions. R.S., J.W., U.A.H., M.K., P.L., and H.S. participated in study design and analyses of data. All authors discussed the results and commented on the manuscript.

## Ethics Declaration

I.H. is the President and Founder of Rakta Therapeutics Inc., a company involved in the development of heme transporter-related diagnostics.

## Supplemental data figures and tables

**Supplemental Data Figure 1. Mapping and identification of suppressor mutants of the heme exporter MRP-5 in *C. elegans*.**

(A) Immunoblot of protein lysates from transiently transfected HEK293 cells and probed with anti-HA, anti-FLAG, and anti-actin antibodies. (B) Cell pellets of HEK293 cells transiently overexpressing the indicated plasmids, after being treated with HD+SA+10 µM heme for 16h. Peroxidase reporter assays in HEK293 cells transfected with (C) cytosol-APX, (D) mitochondria-APX, and (E) ER-HRP reporters. Transfected cells were supplemented with heme depleted (HD) serum + succinyl acetone (SA) and HD + SA + 1 µM heme (n=3). Error bars represent mean ± SEM. (F) HRG-1 mediated heme uptake over time. HEK293 cells transfected with empty vector or *C. elegans* HRG-1 (CeHRG-1) and treated with 5 µM [^55^Fe]heme for 30 h. Cells were harvested and lysed at different time points, and the amount of radioactivity was measured as counts per minute (CPM), normalized to the protein concentration of each sample. Error bars represent mean ± SEM from 2 biological replicates *p<0.05, **p<0.01, ***p<0.001, ****p<0.0001 (one-way ANOVA with Dunnett’s multiple comparison test) (G) Representative immunofluorescence images show that MRP-5 localizes to plasma membrane and trans-Golgi compartments in HeLa cells. (H) Representative confocal image of MRP-5-GFP expression in a whole worm. Scale bar as indicated. (I) Representative images of WT, *mrp-5 (ok2067)*, and suppressor strain *(IQ1005).* Synchronized P_0_ worms were seeded on NGM plates with and without heme and allowed to lay F_1_ progeny. Images were taken 3 days after seeding P_0_ worms. Scale bar, 1 mm. (J) Indicated *mrp-5* suppressor mutants alter ZnMP accumulation within the intestine. Scale bar, 50 µm. (K) Gallium Protoporphyrin IX (GaPPIX) assay shows that suppressor mutants restored intestinal heme efflux (n=3). (L) Pre-culturing parental worms (P_0_) under different heme concentrations caused no change in the heme-dependent growth profile of *mrp-5(ok2067)* worms (n=3). (M) Summary of mapping intervals for *mrp-5* suppressors. ll mapping plots were generated using “CloudMap Hawaiian Variant Mapping with WGS and Variant calling workflow” on the Galaxy Web server. For a given chromosome (linkage group, LG), the top plot represents the ratio of Hawaiian to N2 versions of a single nucleotide polymorphism (SNP), where each black dot represents an individual SNP. The red line represents a LOESS regression line of the frequency of Hawaiian SNPs as a function of genetic locus. The bottom plot represents the frequency of pure N2 alleles over a given interval. Red bars represent bin size of 1Mb. Table shows candidate genes generated by compiling all genes containing missense, nonsense, frameshift, or splicing variants (predicted by snpEff function) within the indicated mapping intervals for each suppressor mutant. (N) RNAi of candidate genes in *mrp-5(ok2067)* (n=3). (O) Candidate testing in WT worms by RNAi knockdown (n=3). (P) Candidate testing in WT (N2) worms by combinatorial RNAi against *mrp-5* (n=3). (H) AP-3 subunit suppressor alleles. (Q) The *aps-3(ih1003)* allele contains a substitution at nucleotide position 383 from C to T. This is predicted to result in a nonsense mutation of codon 115 from R to a stop, and result in a C-terminal truncation of 79 amino acids. The *apd-3(ih1004)* allele contains a substitution at nucleotide position 7,853 from C to T. This is predicted to result in a nonsense mutation of codon 761 from Q to a stop codon, and result in a C-terminal truncation of 492 amino acids. The *apm-3(ih1005)* allele contains a substitution at nucleotide position 1968 from G to A. The mutated guanine is predicted to function as a 5’ nucleophilic splice donor for the excision of intron 9. (R) Depletion of AP-3 subunits in the intestine is sufficient to fully restore viability in *mrp-5* mutants (n=6). (S) Loss of AP-3 subunit does not suppress lethality associated with copper transporter *cua-1*. RNAi performed under copper depleted (Bathocuproinedisulfonic acid disodium salt-BCS) and supplemented (CuCl_2_) conditions (n=3). (T) Heme-dependent growth defect and embryonic lethality of *mrp-5(ok2067)* worms can be rescued by chemically disrupting vesicular trafficking. Worms were bleached, and 100 synchronized L_1_ progeny were grown in mCeHR-2 media supplemented with different concentrations of heme and 80 µM dynasore or 100 µM chloroquine for nine days and quantified (worms/ml) by microscopy (n=2). Peroxidase reporter activity analysis of synchronized L_1_s of (U) *P_dpy-7_::ER-HRP::mCherry* (V) *P_unc-119_::ER-HRP::mCherry* and (W) *P_vha-6_::cyto-APX::EGFP* seeded on indicated RNAi plates, harvested at young adult stage, lysed, and then subjected for activity measurements using o-dianisidine and H_2_O_2_ as substrates (n=3). Error bars represent mean ± SEM. *p<0.05, **p<0.01, ***p<0.001, ****p<0.0001 (one-way ANOVA with Dunnett’s multiple comparison test).

**Supplemental Data Figure 2. Human *SLC49A* homologs act as alternate transporters in the suppressor strain.**

(A) RNA-Seq analysis reveals upregulation of heme responsive genes (HRGs) in the *mrp-5* mutants, which are restored in *apb-3;mrp-5*. (B) Venn diagram showing overlap of the number of candidate genes from RNA-Seq data (green) that contain more than one transmembrane domain (pink). (C) Phylogenetic analysis of *SLC49A* homologs in *C. elegans* and humans. Protein sequences were aligned and phylogenetic tree was generated using the neighbour-joining method in MEGA11. The tree is drawn to scale. The evolutionary distances are in the units of the number of amino acid substitutions per site. (D) Heat map of the RNA-Seq analysis shows regulation of the *SLC49A* homologs. (E) Fragments per kilobase of exon per million mapped fragments (FPKM) values of *SLC49A* homologs in WT worms (n=3). Lethality assays in (F) synchronized N2 (WT) and (G) *apb-3;mrp-5* worms show *B0416.5* and *C27A7.6* are essential in the suppressor (n=3). Heme dose response assays in (H) *mrp-5(ok2067)* and (I) quadruple knockout (*hrg-1;hrg-4,-5,-6*) mutants after RNAi depleting *B0416.5* or *C27A7.6* (n=3). Choline supplementation does not rescue lethality associated with loss of *B0416.5* or *C27A7.6* in (J) N2, (K) *apb-3;mrp-5*, (L) *mrp-5(ok2067)* and (M) *hrg-1;hrg-4,-5,-6* worms (n=3). Error bars represent mean ± SEM. *p<0.05, **p<0.01, ***p<0.001 (one-way ANOVA with Dunnett’s multiple comparison test). (N). Representative confocal image of a whole worm showing B0416.5-GFP expression. (O) Representative confocal image of a whole worm showing C27A7.6-GFP expression. Scale bars as indicated. (P) Representative images and quantification of ZnMP assay in wildtype (WT), *hrg-13*, and *hrg-14* KO worm strains. Scale bar, 10 µm. Error bars represent mean ± SEM from 2 biological replicates. *p<0.05, **p<0.01, ***p<0.001 (one-way ANOVA with Dunnett’s multiple comparison test).

**Supplemental Data Figure 3. Conserved role of SLC49A family transporters in vertebrate erythropoiesis.**

(A) Heme binding sites in SLC49A3 (top, blue) and CeHRG-14 (bottom, light purple) share similar physicochemical characteristics. The heme propionate groups are hydrogen bonded to hydroxyl-groups of amino acid sidechains, the heme plane is sandwiched between two aromatic amino acids (Phe 355/382 and Trp65/295, respectively) and the hydrophobic vinyl face of the protoporphyrin ring is embedded in a cage of hydrophobic amino acids. (B) Molecular dynamics simulations of human SLC49A3. *Top panel* shows backbone RMSD values for the full apo SLC49A3 (blue curves) or the transmembrane core comprising residues 25-468 (red curves) during 200 ns MD simulations compared to the starting structure. Stable conformations are reached after about 50 ns. The longer equilibration times and increased fluctuations of the full transporter compared to the core can be attributed to flexible loops and N- and C-termini. The other two panels show backbone RMSD values for MD simulations of inward (*middle panel*) and outward (*bottom panel*) facing heme-bound SLC49A3. (C) Molecular dynamics simulations of *C. elegans* HRG-14. *Top panel* shows backbone RMSD values for the full apo *Ce*HRG14 (blue curves) or the transmembrane core, comprising residues 43-468 (red curves), compared to the starting structure. A stable conformation is reached readily, with the protein core showing less fluctuation over time due to the absence of flexible loops and disordered N- and C-termini. *Bottom panel* shows backbone RMSD values HRG-14 in complex with heme in the outward open state. (D) Heme is stably bound in MD simulations of *hs*SLC49A3 and *Ce*HRG-14. Top panel represents RMSD of the heme-Fe atom after least square fit of the transporter backbone showing stable heme-coordination for heme-bound to the inward facing SLC49A3. *Middle and bottom panels* represent RMSD of Fe atom for heme-bound to outward facing SLC49A3 and HRG-14 respectively. (E) Schematic showing deletion of human SLC49A3 in K562 cells. (F) WT and SLC49A3 KO cells were treated twice with different concentrations of sodium butyrate (NaB) and harvested 72h after second treatment. Cells were stained with benzidine, as an indirect method to quantify hemoglobinization. (G) Percentage of dead cells in WT and SLC49A3 KO cells with and without NaB treatment, when stained with viability dyes Annexin V and 7AAD. (H) WT and SLC49A3 KO cells were treated twice with different concentrations of hemin and harvested 72h after second treatment. Cells were stained with benzidine, as an indirect method to quantify hemoglobinization. (I) Percentage of dead cells in WT and SLC49A3 KO cells with and without NaB treatment, when stained with viability dyes Annexin V and 7AAD. (J) Phylogenetic analysis of *SLC49A* homologs in zebrafish and humans. Protein sequences were aligned and phylogenetic tree was generated using the neighbour-joining method in MEGA11. The tree is drawn to scale. The evolutionary distances are in the units of the number of amino acid substitutions per site. Alpha-Fold 3 structure predictions of zebrafish homologs (K) SLC49A2a (L) SLC49A2b and (M) SLC49A4. The N and C domains are colored in darker and lighter shades respectively. ChimeraX 1.9 has been used for rendering models. (N) Representative images of 48 hpf zebrafish embryos injected with 0.8ng, 1.6ng, or 3.2ng of *slc49a4* morpholino and stained with *o*-dianisidine to visualize RBCs. Left panels: stained embryos showing RBC distribution. Right panels: corresponding ImagePro quantification with blue overlays indicating gated regions. (O) Quantification of RBC counts show a significant dose-dependent reduction in *slc49a4* morpholino-injected embryos compared with controls. Sample sizes: uninjected (n = 84), standard control 1.6ng (n = 57), *slc49a4* MO 0.8ng (n = 60), 1.6ng (n = 60), and 3.2ng (n = 50). Error bars represent mean ± SEM. *p<0.05, **p<0.01, ***p<0.001, ****p<0.0001 (one-way ANOVA with Dunnett’s multiple comparison test).

**Supplemental Data Figure 4. Biochemical characterization of HRG-13 and HRG-14 in mammalian cells.**

(A) Representative image showing pellets of HEK293 cells transiently expressing the indicated plasmids when treated with HD+SA+10 µM heme. Peroxidase reporter assays in HEK293 cells transfected with (B) ER-HRP, (C) cytosol-APX, and (D) mitochondria-APX reporters under HD+SA and HD+SA+1 µM heme conditions. Error bars represent mean ± SEM from 3 biological replicates. [^55^Fe]heme measurements in (F) cell lysates and (G) media in HEK293 cells transiently transfected with the indicated constructs. Error bars represent mean ± SEM from 2 biological replicates. *p<0.05, **p<0.01, ***p<0.001 (one-way ANOVA with Dunnett’s multiple comparison test).

**Supplemental Data Video 1.** MO injections into Tg(gata1:GFP) embryos decreased gata1 erythroid cells confirming that loss of slc49a4 compromises erythropoiesis in vivo (related to Fig 4M).

**Supplemental Data Video 2.** Live cell imaging video shows vesicular coalescence between heme importer CeHRG-1 (red) and exporter HRG-13 (green) under heme depleted (HD+SA) conditions (related to Fig 5J).

**Supplemental Data Video 3.** Live cell imaging video shows vesicular coalescence between heme importer CeHRG-1 (red) and exporter HRG-13 (green) in the presence of 50 µM added heme (related to Fig 5J).

**Supplemental Data Video 4.** Live cell imaging video shows vesicular coalescence between heme importer CeHRG-1 (red) and exporter HRG-14 (green) under heme depleted (HD+SA) conditions (related to Fig 5K).

**Supplemental Data Video 5.** Live cell imaging video shows vesicular coalescence between heme importer CeHRG-1 (red) and exporter HRG-14 (green) in the presence of 50 µM added heme (related to Fig 5K).

## Methods

### *C. elegans* culture and maintenance

*C. elegans* were maintained in axenic cultures, where worms were grown in continuously shaken liquid mCeHR2 medium ^50^. Alternatively, worms were cultured on nematode growth medium (NGM) plates seeded with either OP50 or HT115(DE3) bacteria at 20°C. The composition of the NGM plates is provided for reference. Routine maintenance procedures, synchronization of worm lifecycles, crosses, and general observations followed established methods described by Epstein and Shakes ^51^

### Synchronization of worms

For harvesting worms on NGM plates, M9 buffer was used to wash them off the agar and remove any bacteria, repeated three times. Worms grown in liquid mCeHR-2 medium were harvested by centrifugation at low speed (800g) for 5 minutes ^50^. After removing the supernatant, the worms were resuspended in a salt solution (0.1 M NaCl) and allowed to settle on ice for 5 minutes. Supernatant was then aspirated. Gravid worms were exposed to a solution of sodium hypochlorite and sodium hydroxide to release the embryos. The mixture was then centrifuged again at 800g for 1 minute. After removing the supernatant, the embryos were washed twice with sterile water and resuspended in M9 buffer for overnight hatching.

### Worm strains

*B0416.5-GFP (PHX8897)* and *C27A7.6-GFP (PHX8895*) worm strains were generated using CRISPR-Cas9 by Suny Biotech. *MRP-5-GFP (IQ1801*) was generated using nested CRISPR ^52^. crRNA synthesis was done using guide RNA design tool from Integrated DNA Technologies (IDT) and ordered from Dharmacon, along with tracrRNA. crRNA used for first round was 5’ ATATCGAAGTTGTTCAGTGA 3’, while the second round was 5’ ACTGGAGTTGTCCCAATCGA 3’. Repair template used for first round of injection: 5’ TCTGAAGTAAAAGAGACGAGCAGTGACACAGATATTGAGGTGGTGCAAGGAGCATCGGGAGCCTCAGGAGCATCGAGTAAAGGAGAAGAACTTTTCACTGGAGTTGTCCCAATCGATGGAGACTATAAGGATCATGATATTGATTATAAAGACGATGATGACAAGTGAAGGATTTCCCATATTTGCCATTATTAAATTTT 3’. Repair template used for second round 5’ GGAGCATCGGGAGCCTCAGGAGCATCGAGTAAAGGAGAAGAACTTTTCTCTTGTTGAATTAGATGGTGATGTTAATGGGCACAAATTTTCTGTCAGTGGAGAGGGTGAAGGTGATGCAACATACGGAAAACTTACCCTTAAATTTATTTGCACTACTGGAAAACTACCTGTTCCATGGGTAAGTTTAAACATATATATACTAACTAACCCTGATTATTTAAATTTTCAGCCAACACTTGTCACTACTTTCTGTTATGGTGTTCAATGCTTCTCGAGATACCCAGATCATATGAAACGGCATGACTTTTTCAAGAGTGCCATGCCCGAAGGTTATGTACAGGAAAGAACTATATTTTTCAAAGATGACGGGAACTACAAGACACGTAAGTTTAAACAGTTCGGTACTAACTAACCATACATATTTAAATTTTCAGGTGCTGAAGTCAAGTTTGAAGGTGATACCCTTGTTAATAGAATCGAGTTAAAAGGTATTGATTTTAAAGAAGATGGAAACATTCTTGGACACAAATTGGAATACAACTTCAACTCACACAATGTATACATCATGGCAGACAAACAAAAGAATGGAATCAAAGCTGTAAGTTTAAACATGATTTTACTAACTAACTAATCTGATTTAAATTTTCAGAACTTCAAAATTAGACACAACATTGAAGATGGAAGCGTTCAACTAGCAGACCATTATCAACAAAATACTCCAATTGGCGATGGCCCTGTCCTTTTACCAGACAACCATTACCTGACCACACAATCTGCCCTTTCGAAAGATCCCAACGAAAAGAGAGACCACATGGTCCTTGTTGAGTTTGTAACAGCTGCTGGGATTACACATGGCATGGATGAACTATACAAAGACTACAAAGATCACGATGGAGACTATAAGGATCATGATATTGATTATAAAGACGATGATGACAA 3’. The repair templates were amplified from pDD282 (plasmid kindly donated by Dr. Xiaofei Bai). *hrg-5,hrg-4,hrg-6* was generated using CRISPR/Cas9. crRNAs used were: 5’ TCCTAGATGTTAAGTTGTGGAGG 3’ and 5’ GAAGTCAATTATTCACCCAGAGG 3’. Repair template used: 5’CTAGAACAATCCTAATAATCCTAGATGTTAAGTTGTGAGCCAGATGATCCATCAACTTGGAAGTTTTAGATG 3’. All the worm strains used in this study have been enlisted in **Supplemental Data Table 1**.

### Generation of knockout worm strains

The *c27a7.6* knockout worm was generated by CRISPR/Cas9-mediated genome editing. Two guide RNAs (AAGGATACTTCGGCAAATGA and TACAGCTCGAACGGCTGCGT) targeting the second and last exons of *c27a7.6* were inserted into a modified pDD162 vector ^53,54^. The gRNA plasmids and a *dpy-10* co-CRISPR plasmid ^55^ were injected into the gonads of adult N2 worms. Dumpy progeny were isolated and screened for mutation in *c27a7.6* by PCR with primers GCATTCCTCACAAAAACAGAGATGCAG, GTTCAGTCATTACATTGAGAATGGTGAG, and GGTGCAAATGAACTGGTGAAGTCTAG. The deletion was verified by sequencing, and the mutant was backcrossed to the N2 strain four times.

The *b0416.5* mutant worm was generated by inserting a *myo-2p::gfp* transgene into its first exon following a previously reported method of CRISPR/Cas9-based homologous recombination ^56^. A gRNA (TGGATTGGATACGCCCCGTC) targeting the first exon of *b0416.5* was inserted into the modified pDD162 (Dickinson et al., 2013; Kang et al., 2020). A repair template was constructed by cloning the sequences flanking the gRNA cutting site (666-bp upstream and 668-bp downstream) into the pDD317 vector (Addgene). The gRNA plasmid and the repair template were injected into adult N2 worms. The progeny were cultured in the presence of hygromycin, and the survived roller worms with pharyngeal GFP expression were selected for genotyping with primers GATGACCTGGAAACAAAGAGACCC, GCATAGTGAAAGTATCCCTCAATGTCTC, and CTCTCCATAGAAGCTATTGACGTAG. The self-excising cassette was removed by culturing the mutant worms at 34℃ for 4 hours. Non-roller mutants were selected for verification of the insertion by sequencing. The mutant was backcrossed to the N2 strain four times.

### Ethyl Methanesulfonate (EMS) mutagenesis screen

Adapted from ^57^. Synchronized worms were grown axenically in mCeHR-2 media in a T-25 flask. Once worms reached the early L4 stage, media was moved from the flask to a 15 mL conical tube. The worm suspension was washed three times (one wash consists of spinning at 800 g for 5 mins, followed by resuspension in 10 mL M9). For the final wash, the worm suspension was resuspended in 2 mL M9. In a laminar flow hood, 0.1 M EMS was made by adding 20µL liquid EMS (Sigma #M-0880) to 2 mL M9 buffer. The solution was gently inverted until EMS dissolved. The 2 mL worm suspension and 2 mL 0.1M EMS was combined for a final volume of 4 mL of 50 mM EMS and worm suspension. The 15 mL conical was closed loosely to allow gas exchange and put inside a container in a 20°C incubator rocking at 70 RPM for 4 h. Following 4h 50 mM EMS treatment, worms were washed twice with M9 as described above and moved back to a T-25 flask containing mCeHR-2 and allowed to grow. All EMS contaminated material (pipette tips, tubes, gloves, etc.) was soaked in EMS inactivating solution (0.1 M NaOH, 20% w/v Na_2_S_2_O_3_) for at least 24 h, and all EMS solution were mixed with equal volume EMS inactivating solution for 24h.

### RNA interference by feeding

RNAi was performed by feeding worms bacteria engineered to express double-stranded RNA (dsRNA) complementary to the target gene. For RNAi feeding all clones were obtained from the established Ahringer library. RNAi bacteria was grown and spotted onto NGM plates containing antibiotics (50 μg/mL carbenicillin and 12 μg/mL tetracycline) and 2mM isopropyl-β-D-thiogalactopyranoside (IPTG) to induce dsRNA production. L_1_ stage larvae were then placed on these RNAi plates and incubated at the desired temperature (20°C) for 72h. For combinatorial RNAi, equal amounts of each bacteria (as measured by OD600) was combined. For heme supplementation, LB used for bacteria culture was directly supplemented with the required hemin chloride concentration, and then shaken at 37°C.

### Hemin chloride/ Sodium butyrate preparation

To prepare a 10 mM hemin chloride solution, 130 mg of hemin chloride powder (Frontier CAS# 16009-13-5) was added in 15 mL of 0.3 M NH_4_OH. The pH of the solution was then adjusted to 8.00 (+/- 0.05) using concentrated HCl. Finally, the solution was brought to a final volume of 20 mL with an additional 0.3 M NH_4_OH (pH 8.0). A 150mM sodium butyrate (Sigma-Aldrich, #B5887) stock solution was dissolved in 1× DPBS freshly for each experiment. Both the solutions were sterilized by passing through a 0.22 μm filter before use.

### Worm growth assay

Approximately 100 bleached and synchronized L_1_ worms were grown axenically in a 12-well plate under different heme concentrations, and the worms were finally counted after 9 days to estimate the total number of worms/ml of media.

### Mapping cross experiment

Crossing plates consisted of 35 mm NGM plates with a 10 µL spot of OP50 bacteria seeded in the center of the plate. Heme supplemented crossing plates used both 200µM hemin chloride added in the NGM agar, as well as in the LB OP50 culture. The P_0_ generation consisted of fifteen Hawaiian *mrp-5(ok2067/o)* males and one late L4 suppressor mutant hermaphrodite. The P_0_ were placed on a heme supplemented crossing plate. The inclusion of an *mrp-5(+/+)* and *mrp5(ok2067/ok2067)* (non-suppressor) were critical controls for downstream screening of F_2_ recombinant lines for *mrp-5* suppression. Once F_1_ progeny were laid, P_0_ were picked off and genotyped to confirm their *mrp-5* allele, and their respective Hawaiian/N2 backgrounds. Each F_1_ animal was picked clonally to 24-well NGM plates supplemented with 200µM hemin chloride in the NGM. F_1_ animals were observed, and the frequency of F_1_ males was used as a relative gauge of mating success. Once F_1_ hermaphrodites laid F_2_ progeny, F_1_ were genotyped for N2 and Hawaiian backgrounds as described. Bona fide cross progeny (heterozygous for Hawaiian and N2 background) were identified, and ∼300 F_2_ from true F_1_ cross progeny were picked out clonally to 24-well NGM plates without heme supplementation. F_2_ animals were allowed to grow and subsequently wells were screened for *mrp-5* suppression, indicated by hatching/growing F_3_ animals. Around 50 recombinant F_2_ lines and their F_3_ progeny were pooled for sequencing.

### Genomic DNA extraction

Worms were collected from plates with M9; all pipetting of worms was done with glass Pasteur pipettes. Collected worm pellets were washed with 10x volume of TE five times to remove bacteria and aspirated to a final volume of 100 µL. The pellet was sonicated with a Bioruptor on high setting, 30s on, 30s off for two 15-minute cycles. Next, 50 µL of Proteinase K (10 mg/mL) was added followed by incubation at 65°C for one hour, until the solution was clear. Next, 20 µL of RNaseA (10 mg/mL) was added and then the solution was incubated at 37°C for 30 mins. Following incubation, the DNA purified with the Qiagen PCR purification kit and eluted with 30 µL TE. Quantification of DNA was done with PicoGreen (ThermoFisher).

### Library preparation

Library construction was done with the NuGen Ovation SP^+^ Ultralow DR kit, using 50 ng sheared genomic DNA samples. Library integrity was checked by Agilent High Sensitivity DNA Assay Kit.

### Sequencing

The sequencing was at the NIDDK Genomics Core on an Illumina HiSeq 2500; at least 20x genome coverage (read depth) was obtained for each sample to ensure accurate variant calling. In addition to the three suppressors analyzed, the two parental strains were also sequenced (N2; *mrp5(ok2067/ok2067*) pre-mutagenesis strain to allow identification of novel EMS induced mutations, as well as Hw; *mrp-5(ok2067/ok2067)* background which contains the paternal contribution for the mapping cross).

### Sequence analysis

All downstream analyses of the gripped FASTQ files were performed using CloudMap80. First, FASTQ files were concatenated for each sample sequenced. Variants present in either parental strains [Hawaiian; *mrp-5(ok2067/ok2067)* (paternal strain) or N2; *mrp-5(ok2067/ok2067*) (pre-mutagenesis strain)] were subtracted using “CloudMap Subtract Variants workflow (1 set candidates, 2 sets variants to subtract)” workflow published by gm212380 to constrain the set of variants to those induced by EMS. Using the output from this workflow, all other 24 analysis was conducted using the “CloudMap Hawaiian Variant Mapping with WGS and Variant Calling workflow” published by gm2123. ^58^

### Embryonic lethality assay

Worms were synchronized and placed on RNAi plates (NGM plates with Noble Agar (Difco CAS 214320)) with bacterial cultures grown in 1/4 Luria-Bertani broth (LB) (2.5 % tryptone, 1.25 % yeast extract, and 10 % NaCl weight by volume, 50 μg/mL carbenicillin, 12 μg/mL tetracycline, and 2 mM IPTG) with no added heme or different concentrations of added heme. After 48 h, young adult worms were clonally picked to fresh RNAi plates and allowed to lay embryos for 24 h. After another 24 h, these worms were then moved to new RNAi plates and hatched worms and unhatched embryos were counted from the previous plate. For the copper supplementation experiment, BCS and CuCl_2_ were added to the NGM plates, which were then seeded with the RNAi bacteria. For choline supplementation, 100 μM choline chloride was added to the NGM plates, after which RNAi was performed.

### Rescue *mrp-5(ok2067)* worms with trafficking inhibitors

P_0_ generation of *mrp-5 (ok2067)* worms were maintained in the mCeHR2 medium supplemented with 200µM heme and synchronized as described above. After 24h, approximately 100 synchronized L_1_ worms were placed into 1mL mCeHR2 medium containing heme and trafficking inhibitors in each well of 24-well plates (day 0). The total number of worms in every well was counted on day 9 under a dissection microscope (Leica). Dynasore (Santa Cruz Bio, Cat#sc-202592) was prepared as 80mM stock in DMSO. Chloroquine (MP Biomedicals, Cat#193919) was prepared as 100mM stock in sterile water.

### Immunoblotting

Worms were harvested, lysed (using Beadbeater) and total protein concentration was measured by Bradford’s assay, as described previously.^59^ Mammalian cells were lysed using standard lysis protocol ^60^. Membranes were probed overnight in 4°C in blocking buffer containing primary antibody of interest: mouse anti-GFP (Biolegend CAS 902603), rabbit anti-HA (Sigma, #H7908), mouse anti-actin (DSHB, #JLA20), or anti-tubulin (DSHB, #AA4.3) at a 1:1000 concentration. Blots were developed using enhanced chemiluminescence using Azure imaging systems. Quantification of blots was done using ImageJ, where tubulin was used as a loading control.

### Heme analog uptake assays

Zinc mesoporphyrin (ZnMP) and gallium protoporphyrin IX (GaPPIX) were prepared as 10 mM stock solutions, dissolved in NH_4_OH (pH8). For ZnMP uptake assay, L_4_ worms were harvested and incubated in 10 μM ZnMP for 16h, washed with M9 and then imaged, as outlined previously ^59^. For GaPPIX toxicity assay, L_4_ worms were added to NGM plates containing different concentrations of GaPPIX and seeded with OP50 bacteria. After 48h, worms were scored as dead, if they were unresponsive to physical stimulus.

### LysoTracker staining

50 μM LysoTracker (LysoTracker Blue DND-22 CAS L7525) solution was prepared in M9. Worms were incubated in this solution for 15 mins at 20°C in the dark. They were then washed and allowed to de-stain for another one hour at 20°C in the dark, followed by imaging.

### Worm microscopy

Representative confocal worm images were taken using Zeiss LSM710 and Nikon A1 with a 63X oil immersion objective. After worms were harvested, they were added to slides with 2% agar pads. Worms were immobilized using 10 mM Levamisole (MP Bio, CAS 155228), and then mounted on cover slips before visualization. All worm images have been denoised using Nikon NIS-elements software.

### RNA-Seq analysis

Synchronized L1 larvae of IQ1904 (WT brood-mate), IQ1903 (*mrp-5(ok2067)*), IQ1902 (*apb-3(ok429)*), and IQ1901 (*mrp-5(ok2067);apb-3(ok429*)) strains were grown on 10 cm NGM plates supplemented with OP50 bacteria and 10 µM heme. Worms were collected at late L_4_ stage in M9 buffer, followed by three additional washes with M9 buffer. Total RNA was extracted with RNeasy kit (Qiagen, Germany). A total of 12 samples were sequenced using Illumina’s HiSeq-2500, with triplicate of each genotype. Single-end 50 base reads were sequenced and checked by FastQC, version 0.11.5. The reads were aligned to *C. elegans* UCSC reference genome ce10 using STAR, version 2.5.2b. Differentially expressed genes (DEGs) were identified using DESeq2, version 1.40.2 and R version 4.3.1. Log2 fold changes of gene expression was shrunk by the ‘lfcShrink (ashr)’ function of DESeq2. The False Discovery Rate (FDR) was used to determine the statistical significance with the cutoff value (adjusted p-value) of 0.05 set for DEGs. All the sequencing data including read counts per gene will be deposited to NCBI Sequence Read Archive (SRA). Transmembrane domain analysis was performed using TMHMMM-2.0 (https://services.healthtech.dtu.dk/services/TMHMM-2.0/) Gene Ontology (GO) analysis was conducted using WormBase Gene Set Enrichment Analysis tool, and results were plotted using R ^61,62^.

### DNA cloning

Worm B0416.5 and C27A7.6 ORFs were synthesized using TWIST. HA epitope tag was added by PCR and then this construct was digested with XhoI and BamHI. This was then ligated into pEGFP-N1 cut with XhoI and BamHI. Human codon optimized worm MRP-5, tagged with HA, was synthesized using TWIST. MRP-5-HA was ligated into pcDNA3.1 for mammalian cell expression.

### Integration of extrachromosomal arrays

Extrachromosomal transgenic array *Pvha-6::HRG-14-HA-mScarlet::unc-54 3’UTR* was integrated using trimethylpsoralen (TMP)-mediated UV crosslinking to promote chromosomal incorporation, following established procedures for array integration. Briefly, animals carrying the desired extrachromosomal array were exposed to TMP followed by UV irradiation, and surviving progeny were recovered and screened for stable inheritance of the fluorescent transgene. Candidate integrated lines were identified based on *mScarlet* expression across multiple generations. Integrated line was established and subsequently backcrossed to the wild-type background four times prior to experimental use.

### Immunoprecipitation of *C*. *elegans*

Ce-HRG1 IP eluates were subjected to reduction with 5 mM tris(2-carboxyethyl)phosphine (TCEP) and subsequent alkylation using 10 mM iodoacetamide (IAA). Reduced and alkylated proteins were cleaned via the protein aggregation capture (PAC) method ^63^, after which they were enzymatically digested overnight at 37°C using Lys-C and trypsin proteases. The resulting peptide mixtures were dried prior to LC–MS/MS analysis.

The beads obtained after immunoprecipitation of MRP-5 were resuspended in 2M Urea, 100mM Tris-Cl, pH 8.0 followed by reduction and alkylation as mentioned above. Reduced and alkylated bead-bound proteins were digested with lysC and trypsin at 37°C overnight. The digestion was quenched by addition of formic acid to a final concentration of 5%. The digested peptides were desalted using SP3-based peptide clean-up protocol ^64^. The resulting peptides were dried and analyzed by LC-MS/MS.

Dried tryptic peptides were reconstituted in 5% formic acid and injected onto a PepSep C18 reverse-phase column (150 mm × 150 µm, 1.7 µm particle size) maintained at 59°C, with electrospray ionization into an Orbitrap Astral mass spectrometer. Chromatographic separation followed a trap-and-elute workflow performed on a Vanquish Neo UHPLC system. The mobile phases consisted of water with 0.1% formic acid (A) and acetonitrile with 0.1% formic acid (B). A 15-minute gradient was applied as follows: 5% B from 0–1 min (2.45 µL/min), 5–15% B from 1–5 min (1.75 µL/min), 15–25% B from 5–12.6 min (1.75 µL/min), 25–38% B from 12.6–13.6 min (1.75 µL/min), and 38–80% B from 13.6–13.7 min (2.45 µL/min), with the column held at 80% B until 15 min (2.45 µL/min).

Data-independent acquisition (DIA) was performed on the Astral mass spectrometer operating in positive electrospray ionization mode. MS1 spectra were collected at 240,000 resolution across an m/z range of 380–980, with a normalized AGC target of 500% and a maximum injection time of 3 ms. DIA scans employed sequential 4 m/z-wide isolation windows covering the same range. MS2 spectra were acquired at 80,000 resolution using a normalized higher-energy collisional dissociation (HCD) energy of 25%, a normalized AGC target of 500%, and a maximum injection time of 7 ms

Thermo RAW data files were processed with DIA-NN against an in silico spectral library generated from the *Caenorhabditis elegans* reference proteome (UniProt ID: UP000001940)^65^. The DIA-NN outputs were subsequently analyzed in FragPipe Analyst to determine differentially regulated proteins ^66^.

### Yeast three-hybrid (Y3H) assay

Y3H vector construction and assays were performed as described previously ^67,68^. Briefly, codon-optimized APM-3, APD-3 and APS-3 were cloned into pACT2, pGADT7 and pBridge vectors. The N- and C-terminal cytoplasmic sequences of HRG-1, HRG-4, MRP-5, HRG-13 and HRG-14 were cloned into pGBT9 and pBridge-APS-3 for Y2H and Y3H assays, respectively. pBridge-APS-3 plasmids carrying N- and C-terminal constructs and pGADT7-APD-3 were co-transformed into HF7c yeast strain and selected on SD glucose agar plates lacking leucine, tryptophan and methionine but containing histidine. The Y3H growth assay was performed on SD glucose -Leu, -Trp, -Met, -His plates containing increasing concentrations of 3-amino-1,2,4-triazole (3-AT).

### Zebrafish lines and maintenance

All zebrafish (*Danio rerio*) experiments were performed in compliance with National Institutes of Health (NIH) guidelines for animal handling and research under an Animal Care and Use Committee–approved protocol. Zebrafish were maintained and bred as described previously ^69^. Embryos were raised at 28.5 °C in an incubator under standard conditions. The following strains were used: TAB5 wild-type and *Tg(gata1:GFP)* ^70^.

### CRISPR/Cas9 crispant and morpholino injections

CRISPR single-guide RNA (sgRNA) target sites were identified using CRISPOR ^71^. sgRNAs were synthesized by Synthego and incubated with EnGen Spy Cas9 NLS protein (New England Biolabs, Ipswich, MA) at 37 °C for 5 min to generate ribonucleoprotein (RNP) complexes. Wild-type embryos were injected at the one-cell stage using a PicoPump microinjector (World Precision Instruments, Sarasota, FL) following standard protocols ^69^.

For Crispant experiments, injection mixes contained two sgRNAs (200 pg each) together with Cas9 protein and were delivered at the one-cell stage. Treatment groups included *sal49A2a* (scl49A2a-E1-T1 and slc49A2a-E2-T2), *slc49A2b* (slc49A2b-E1-T1 and slc49A2b-E2-T2), and *slc49a4* (slc49a4-E1-T1 and slc49a4-E3-T2). To account for potential functional redundancy, a combined treatment group was generated in which one sgRNA targeting *slc49A2a* (slc49A2a-E2-T2) and one sgRNA targeting *slc49A2b* (slc49A2b-E1-T1) were co-injected together with Cas9. Follow-up experiments for *slc49a4* were performed using a single sgRNA (slc49a4-E1-T1) at either 200 pg or 400 pg. A complete list of sgRNAs, target exons, and primers for genotyping is provided in **Supplemental Data Table 5**.

Morpholino antisense oligonucleotide (Gene Tools, Philomath, OR) was designed against *slc49a4* along with a standard control morpholino. Embryos were injected with the *slc49a4* translation-blocking morpholino (5’-CACCCATCGAGAACACACGGTATCA-3’) at doses of 800, 1600, 3200, or 4800 pg, and with the standard control morpholino (5’-CCTCTTACCTCAGTTACAATTTATA-3’) at 1600 and 4800 pg. Morpholino injections were performed in wild-type, and *Tg(gata1:GFP)* transgenic zebrafish using standard protocols.

CRISPR-STAT analysis was performed on Crispant embryos to confirm sgRNA activity ^72^. DNA was extracted and amplified using REDExtract-N-Amp PCR ReadyMix (Sigma-Aldrich, St. Louis, MO) according to the manufacturer’s instructions.

### *o*-dianisidine staining and imaging

Embryos were collected at 48 hpf and incubated in *o*-dianisidine solution (MP Biomedicals, Costa Mesa, CA) for 15 min in the dark ^73^. Samples were fixed in 4% paraformaldehyde (PFA) for 1 h at room temperature and subsequently bleached in 30% hydrogen peroxide for

8 min to remove pigment prior to imaging. Brightfield images of stained embryos and fluorescence imaging of transgenic lines at 3 dpf were acquired using a Leica M205 microscope equipped with a Leica DFC7000G camera, controlled by the Leica Application Suite X Imaging Software Suite version 3.4.1 (Leica Microsystems, Deerfield, IL). For time-lapse imaging, one frame was captured every 1.5 s for 53 s.

### Zebrafish image analysis and quantification

Images were analyzed using ImagePro software (Media Cybernetics, Rockville, MD) to estimate red blood cell (RBC) numbers. For Crispant experiments, counts from lateral and ventral views were combined to yield a total estimated RBC count per embryo. For morpholino experiments, only ventral yolk views were used to estimate RBC numbers. Graphs and statistical analyses were generated in GraphPad Prism (GraphPad Software, San Diego, CA). Statistical significance was determined using Tukey’s multiple comparisons test.

### Mammalian cell culture and transfection

Mammalian cell culture experiments were performed using HEK293, HeLa cells, and K562 cells. These cells were grown in a humidified incubator at 37°C with 5 % CO2. Dulbecco’s modified Eagle medium (DMEM) supplemented with 10 % FBS and 1 % penicillin/streptomycin/glutamine (PSG) was used as the culture medium for HEK293 and HeLa cells. K562 cells were grown in RPMI-1640 medium supplemented with 10% FBS and 1% PSG. HD media was prepared as described in ^19^. Transfection for immunoblotting, reporter assays, immunofluorescence, and imaging, was done using PolyJet (SignaGen, SL100688).

### Generation of SLC49A3-KO K562 cells

CRISPR-Cas9-mediated gene knockout was performed in K562 cells following established genome engineering strategies ^74^. Single guide RNAs (sgRNAs) targeting SLC49A3 were designed and cloned into pSpCas9(BB)-2A-GFP(PX458). SgRNA sequences were Sg1(5’ TGTCCATGGAGCAGATCAAC 3’), Sg4(5’ AGCGTCCAGGTTTATTGACC 3’). K562 cells were electroporated with the CRISPR-Cas9/sgRNA construct, and transfected populations were expanded under standard culture conditions. Clonal cell lines were isolated by fluorescence-activated cell sorting based on GFP signal, and candidate knockout clones were screened by PCR genotyping with forward (5’-CTGTGGCTCAGCTTTGCACCTG-3’) and reverse primers (5’-CTGGTGCTCTGGGAACCACAAG-3’ for exon 3, and 5’-CTGCAGGTGCGCGCTTGTAATTC-3’ for exon 10). Verified knockout lines were maintained and used for subsequent experiments.

K562 cells were then collected, rinsed once with 1× DPBS, and resuspended to a final 100 µL volume. Cell pellets were disrupted using a Bioruptor sonicator at the high-power setting, with 30-second pulses at 30-second intervals for two 15-minute cycles. Following disruption, 50 µL Proteinase K (10 mg/mL) was added and samples were incubated at 65 °C for 1 hour until a clear solution was obtained. Subsequently, 20 µL RNase A (10 mg/mL) was added, and reactions were incubated at 37 °C for 30 minutes. After these steps, DNA was isolated using the Qiagen PCR purification kit and eluted in 30 µL TE buffer. DNA concentration was measured using PicoGreen (ThermoFisher).

### *o*-dianisidine staining (benzidine) of K562 cells

*o*-dianisidine staining was used to visualize hemoglobin within K562 cells from intact embryos or prepared tissue sections, following previously established protocols (13, 18). In brief, K562 cultures were incubated in 200 µL of staining reagent composed of 0.6% (w/v) o-dianisidine, 25% ethanol, 10 mM sodium acetate, and 0.02% H O for 10 minutes under light-protected conditions. Heme mediates the oxidation of o-dianisidine in the presence of hydrogen peroxide, yielding a dark brown precipitate that marks hemoglobin-containing cells. Both total cell numbers and stained cell counts were determined using a hemocytometer.

### Flow cytometry

K562 WT and *SLC49A3* cell apoptosis were measured by flow cytometry. Around one million of cells were washed and stained with 2µL of Annexin-V-Pacific Blue (BioLegend, 640918) and 7-AAD (eBioscience, 00-6993-50). Cells were acquired using LSRII (BD Bioscience). Results were analyzed using Flow-Jo software (Tree Star, Ashland, OR).

### Immunofluorescence of mammalian cells

HeLa cells were transfected with pcDNA3.1-MRP-5-HA alone, and co-transfected with TGN38-mCherry. Two days post transfection, the cells were fixed using 4% paraformaldehyde (PFA), blocked in 5% BSA, and incubated with primary antibody (anti-HA, 1:100), secondary antibody (Alexa-conjugated anti-rat, 1:5000). HeLa cells were transfected with B0416.5-HA-GFP or C27A7.6-HA-GFP by themselves or along with different markers (LAMP1-mCherry, or CeHRG1-mCherry) and then fixed with 4% PFA. After fixation, cells were stained using 5 μg/ml Wheat Germ Agglutinin (WGA), for 10 minutes at room temperature. Cells were then washed with PBS thrice and subsequently mounted using Antifade (ProLong Gold Antifade Reagent, Invitrogen, CAS P36934). Images were taken and processed using Zeiss Airyscan 980 SR and Nikon A1 confocal microscope. Images have been denoised using Nikon NIS-elements software.

### Oxalic acid heme quantification assay

Transfected cells treated HD + 0.5 μM SA+10 μM hemin chloride were harvested and lysed using 2 M oxalic acid solution by vortexing. Protein concentration was determined using Bradford assay. Samples along with porphyrin and heme standards were incubated in heating block at 100°C and room temperature for 1 h to convert heme to porphyrin. Porphyrin fluorescence was then measured at excitation 400nm and emission 660nm.

### Horseradish/ascorbate-peroxidase assay

HEK293 cells were transfected with equal amounts of empty vector, MRP-5, CeHRG-1, B0416.5 and C27A7.6 along with either ER-HRP or cytosol-APX or mitochondria-APX using PolyJet transfection reagent. The following day, the media was changed to heme depleted (HD) media (DMEM with 10 % heme depleted FBS and 1 % PSG), and 0.5 μM succinylacetone (SA), along with different concentrations of hemin chloride. The following day, cells were harvested and lysed using a buffer containing 150 mM NaCl, 50 mM HEPES, and 1 % Trition X, with protease inhibitor. Total protein concentration for each sample was measured using Bradford Assay. Peroxidase activity was measured as described previously^19^. Each sample was normalized to the peroxidase activity of empty vector at that heme concentration, and to the protein concentration. Similar protocol was followed for worm peroxidase assay, where young adult worms from different RNAi conditions were harvested, lysed, and the total protein concentration was measured using Bradford’s Assay.

### Live cell imaging

HeLa cells were transfected with HRG-13-HA-GFP or HRG-14-HA-GFP along with CeHRG-1-mCherry. The following day the media was changed to HD along with 0.5 μM SA. Again, the day after, fresh OptiMEM + SA media was added and then taken for imaging using the Nikon W1 spinning disk. 50 μM hemin chloride was added while the cells were being imaged. Videos were processed and denoised by Nikon NIS-Elements software. Pearson colocalization coefficient was calculated across the entire time span of the video (3 mins) for each condition using the colocalization tool in Imaris software (Bitplane).

### [^55^Fe]-heme uptake

[^55^Fe]-heme was synthesized by incorporating 18mCi/mg ^55^FeCl_3_ (Perkin Elmer) into protoporphyrin IX (Frontier Scientific) as previously outlined ^75^. HEK 293 cells were seeded at a 60% confluency per 12-well plate in duplicates. Next day, cells were transfected with CeHRG-1 or empty vector. Media was changed the next day. The following day (day of the experiment), cells were incubated with 5 μM [^55^Fe]-heme for different durations ranging from 30 minutes to 30 h. After each time point, cells were harvested, washed with PBS containing 5 % BSA, lysed using buffer containing 150 mM NaCl, 50 mM HEPES, and 1%Triton X, and then transferred to a scintillation cocktail (Ecoscint A Scintillation Cocktail, CAS LS-273). Throughout the experiment media contained heme-depleted serum and 0.5 µM succinylacetone (HD+ 0.5 μM SA). Radioactivity was measured using a scintillation counter. Radioactivity was normalized to the total protein concentration for each sample. ^55^Fe uptake was determined by subtracting the CPM values for samples at 4°C from the samples at 37°C.

### [^55^Fe]-heme efflux

HEK 293 cells were seeded at a 60% confluency per 12-well plate in duplicates. The following day, cells were transfected with different DNA constructs (empty vector, CeHRG-1, MRP-5, B0416.5, C27A7.6, CeHRG-1+ MRP-5, CeHRG-1+ B0416.5 and CeHRG-1+ C27A7.6). Media was changed the next day. The following days cells were incubated with different concentrations of [^55^Fe]-heme for 6 h. Cells were then washed twice with PBS containing 5% BSA, and media was changed to fresh OptiMEM without any added heme, and incubated for another 24 h at 37°C. After this, cells were harvested, lysed using buffer containing 150 mM NaCl, 50 mM HEPES, and 1% Triton X and then transferred to a scintillation cocktail (Ecoscint A Scintillation Cocktail, CAS LS-273). The media was also collected and transferred to a scintillation cocktail. Throughout the experiment HD + 0.5 μM SA media was used. Radioactivity was measured using a scintillation counter and was normalized to the total protein concentration for each sample.

### Image analysis and 3D rendering

All image analysis including quantification of signal and measurement of vesicular volume were performed using Imaris software (Bitplane). The surface module was used for quantification. 3D rendering was also done using Imaris (Bitplane).

### Molecular Dynamics (MD) Simulations

To generate starting structures for all atom MD simulations, Alphafold 3 predictions of human SLC49A3 and *C. elegans* HRG-14 with and without heme were generated using Alphafoldserver ^76^. For unbound SLC49A3 and HRG-14, as well as heme-bound HRG-14, all predictions yielded the outward facing state. Thus, the best ranked model was used as the respective starting structure. For heme-bound SLC49A3. Both inward and outward open conformations were predicted and the best ranked models for either conformation were used. MD simulations of the four conformations were carried out in a representative eukaryotic membrane, asymmetrically assembled from phospholipids, sphingomyelin, glucosylceramide and cholesterol **(Supplemental Data Table 4)**. For an initial placement of SLC49A3 and HRG-14 into the membrane, the respective protein was first superimposed to the structure of the 12-trasmembrane helix protein OCT3 in lipid nanodisc ^77^ (PDB:7ZH0) obtained from Orientations of Proteins in Membranes (OPM) database ^78^. Membrane embedding was then validated by generating vacuum electrostatics with PyMOL 2.5. To minimize the system size for all atom simulations, the disordered C-terminus of human SLC49A3 was cleaved at position Alanine 462. To obtain the input files to simulate the respective proteins in the lipid bilayer, each membrane-oriented structure was then parsed to CHARMM-GUI ^79-81^. Using the membrane builder module ^82-89^, the oriented protein was embedded into a membrane bilayer surrounded by water as well as 0.15 mmol/L each sodium and chloride ions. After minimization and equilibration, GROMACS input files ^81^ were generated. MD Simulations were run using GROMACS 2021.2 (120-200 ns, 2 replicates each). Evaluation of Root Mean Squared Deviations and visualization of structures was done using built-in GROMACS modules, PyMOL and Biovia Discovery Studio Client (BIOVIA, Dassault Systèmes, Discovery Studio Client, v21.1.0, San Diego: Dassault Systèmes, 2020). All proteins showed a stable RMSD relative to the respective starting structure after 40-50 ns, with the heme iron undergoing minimal movement after equilibration. Thus, snapshots of human SLC49A3 and *C. elegans* HRG-14 in the heme-bound, outward open state at 120 ns simulations were used to generate figures for overlays and representative heme binding poses.

### Software

MEGA11 was used to make phylogenetic tree. Protein structure modelling was done using AlphaFold3. Rendering was done using ChimeraX 1.9. All data are presented as mean ± the standard error of the mean (SEM). Statistical significance was determined using one-way or two-way ANOVA or unpaired t test, as indicated in figure legends using GraphPad Prism, version 10.00 (GraphPad Software, Inc). Some figures, where denoted, were generated using BioGDP ^90^. Figures of transporters were generated using pyMOL (The PyMOL Molecular Graphics System, Version 2.5).

---

**Supplemental Data Table 1.** *C. elegans* strains list in the study.

| Strain | Genotype | Source |
| --- | --- | --- |
| N2 | Bristol (wild-type) | CGC |
| IQ5951 | *mrp-5(ok2067) X* | 17 |
| IQ1801 | *mrp-5(ih1801[mrp-5::gfpnovo2<sup>3xFlag</sup>]) X*, CRISPR/Cas9 edit | This study |
| IQ1003 | *mrp-5(ok2067) X; aps-3(ih1003[R115\*]) I*, EMS mutagenesis | This study |
| IQ1004 | *mrp-5(ok2067) X; apd-3(ih1004[Q761\*]) II*, EMS mutagenesis | This study |
| IQ1005 | *mrp-5(ok2067) X; apm-3(ih1005) X*, EMS mutagenesis, loss of intron 9 splicing donor site (g->a) | This study |
| IQ6926 | *hrg-1(tm3199) X; hrg-5&hrg-4&hrg-6(ih6925) IV*, CRISPR/Cas9 edit | This study |
| VP303 | *rde-1(ne219) V; kbIs7[nhx-2p::rde-1 + rol-6(su1006)]* | CGC |
| IQ9101 | *ihIs9101 [vha-6p::ER-HRP-mCherry::unc-54 3'UTR + unc-119(+)]; unc-119(ed3) III; lin-15B(n744) X* | This study |
| IQ9102 | *ihIs9102 [dpy-7p::ER-HRP-mCherry::unc-54 3'UTR + unc-119(+)]; unc-119(ed3) III; lin-15B(n744) X* | This study |
| IQ9104 | *ihIs9104 [unc-119p::ER-HRP-mCherry::unc-54 3'UTR + unc-119(+)]; unc-119(ed3) III; lin-15B(n744) X* | This study |
| IQ9006 | *ihIs9006 [vha-6p::egfp-cytoAPX::unc-54 3'UTR + unc-119(+)]; unc-119(ed3) III* | This study |
| RB662 | *apb-3(ok429) I* | CGC |
| IQ1901 | *apb-3(ok429) I; mrp-5(ok2067) X* | This study |
| IQ6111 | *ihIs6111 [hrg-1p::hrg-1::gfp::unc-54 3'UTR + unc-119(+)]; unc-119(ed3) III* | 16 |
| IQ6161 | *ihIs6161 [vha-6p::hrg-1-HA::sl2::mCherry::unc-54 3'UTR + unc-119(+)]; unc-119(ed3) III* | This study |
| IQ6112 | *ihIs6112 [vha-6p::hrg-1::mCherry::unc-54 3'UTR + unc-119(+)]; unc-119(ed3) III* | This study |
| IQ6192 | *ihIs6192 [vha-6p::hhrg-1::mCherry::unc-54 3'UTR + unc-119(+)]; unc-119(ed3) III* | This study |
| IQ2001 | *mrp-5p::GFPNovo2::3xFLAG X; IQ6161 X* | This study |
| IQ5301 | *vha-6p::c27a7.6-HA::mScarlet::unc-54 3'UTR V* | This study |
| IQ5952 | *vha-6p::hrg-1::mCherry::unc-54 3'UTR X; mrp-5(ok2067) X* | This study |
| CCH1001 | *b0416.5(cck1001)X* | This study |
| CCH1004 | *b0416.5(cck1001)X; cck1014[vha-6p::b0416.5::gfp]* | This study |
| CCH1002 | *c27a7.6(cck1002)V* | This study |
| PHX8897 | *b0416.5(syb8897[B0416.5::gfpnovo2<sup>3xFlag</sup>]) X*, CRISPR/Cas9 edit | This study |
| PHX8895 | *c27a7.6(syb8895[C27A7.6::gfpnovo2<sup>3xFlag</sup>]) V*, CRISPR/Cas9 edit | This study |

**Supplemental Data Table 2:** Adaptor Protein subunits knockdown.

| Gene | Name | Observation | Worm strain |
| --- | --- | --- | --- |
| Y71H2B.10 | *apb-1* | No change | MRP-5::GFP |
| F55A12.7 | *apm-1* | No change |  |
| F29G9.3 | *aps-1* | No change |  |
| T20B5.1 | *apa-2* | Partially decreased |  |
| R160.1 | *apm-2* | Decreased |  |
| F02E8.3 | *aps-2* | Decreased |  |
| R11A5.1 | *apb-3* | Decreased |  |
| W09G10.4 | *apd-3* | Decreased |  |
| F53H8.1 | *apm-3* | No change |  |
| Y48G8AL.14 | *aps-3* | Mislocalized |  |
| Y71H2B.10 | *apb-1* | Partially increased | HRG-1::mCherry |
| F55A12.7 | *apm-1* | No change |  |
| F29G9.3 | *aps-1* | No change |  |
| T20B5.1 | *apa-2* | Decreased |  |
| R160.1 | *apm-2* | No change |  |
| F02E8.3 | *aps-2* | Mislocalized/ Clumping |  |
| R11A5.1 | *apb-3* | Decreased |  |
| W09G10.4 | *apd-3* | Partially decreased |  |
| F53H8.1 | *apm-3* | Mislocalized/ Clumping |  |
| Y48G8AL.14 | *aps-3* | No change |  |

**Supplemental Data Table 3:** Trafficking genes knockdown.

| Gene | Name | Observation | Worm strain |
| --- | --- | --- | --- |
| F46F11.5 | *vha-10* | Decreased | MRP-5::GFP |
| Y49A3A.2 | *vha-13* | Increased/Mislocalized |  |
| ZK180.4 | *sar-1* | Decreased |  |
| T20G5.1 | *chc-1* | Decreased | HRG-13::GFP |
| C01G8.5 | *erm-1* | Decreased |  |
| Y54G11A.10 | *lin-7* | Decreased |  |
| C39F7.4 | *rab-1* | Decreased |  |
| ZK637.8 | *unc-32* | Decreased |  |
| F46F11.5 | *vha-10* | Decreased |  |
| T20G5.1 | *chc-1* | Decreased | HRG-14::GFP |
| F38E11.5 | *copb-2* | Increased/Mislocalized |  |
| T14G10.5 | *copg-1* | Increased |  |
| F12F6.6 | *sec-24.1* | Mislocalized |  |

**Supplemental Data Table 4:**
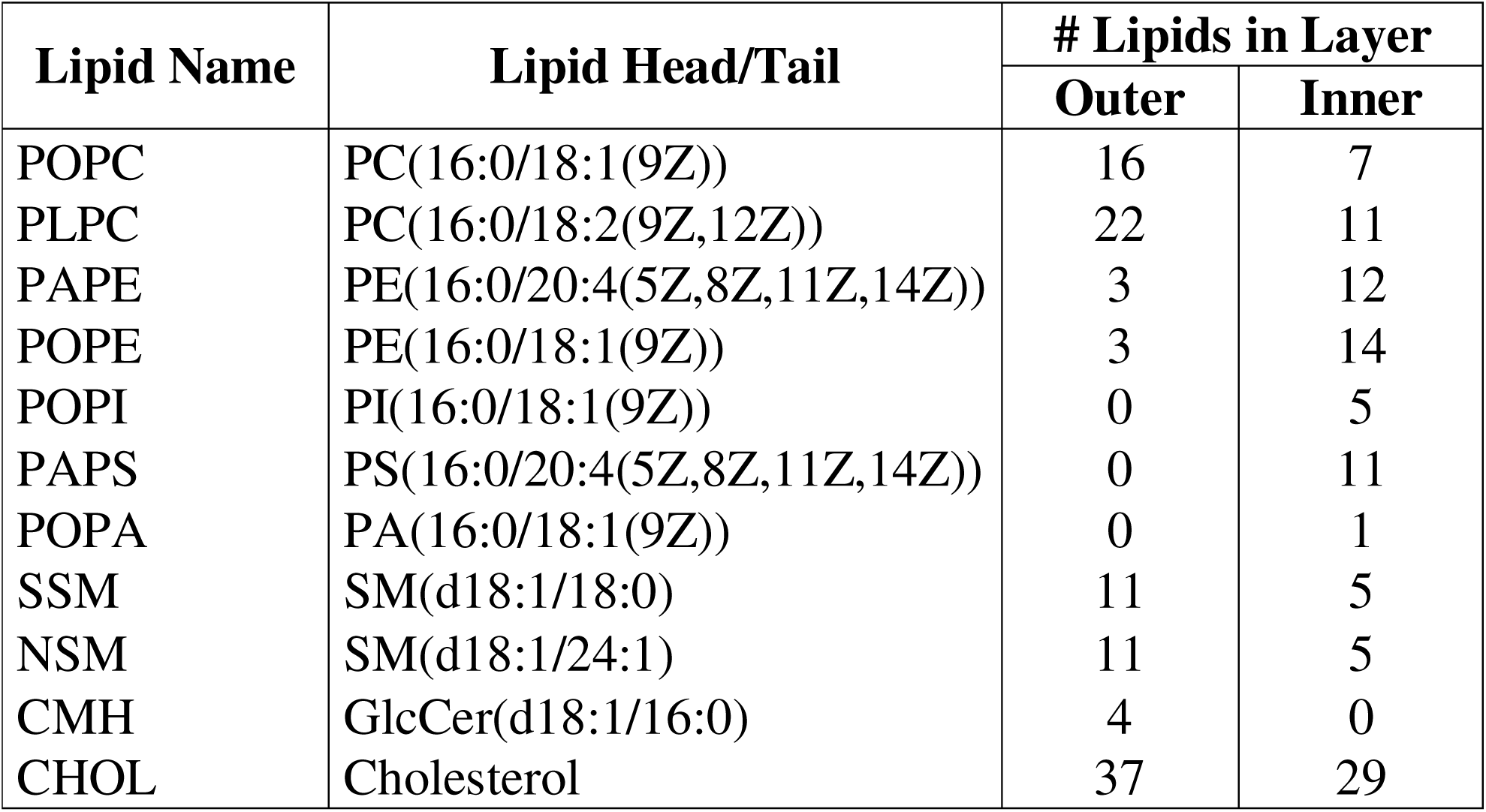
Lipid composition of asymmetric mammalian plasma membrane.

| Lipid Name | Lipid Head/Tail | # Lipids in Layer |  |
| --- | --- | --- | --- |
|  |  | Outer | Inner |
| POPC | PC(16:0/18:1(9Z)) | 16 | 7 |
| PLPC | PC(16:0/18:2(9Z,12Z)) | 22 | 11 |
| PAPE | PE(16:0/20:4(5Z,8Z,11Z,14Z)) | 3 | 12 |
| POPE | PE(16:0/18:1(9Z)) | 3 | 14 |
| POPI | PI(16:0/18:1(9Z)) | 0 | 5 |
| PAPS | PS(16:0/20:4(5Z,8Z,11Z,14Z)) | 0 | 11 |
| POPA | PA(16:0/18:1(9Z)) | 0 | 1 |
| SSM | SM(d18:1/18:0) | 11 | 5 |
| NSM | SM(d18:1/24:1) | 11 | 5 |
| CMH | GlcCer(d18:1/16:0) | 4 | 0 |
| CHOL | Cholesterol | 37 | 29 |

**Supplemental Data Table 5:**
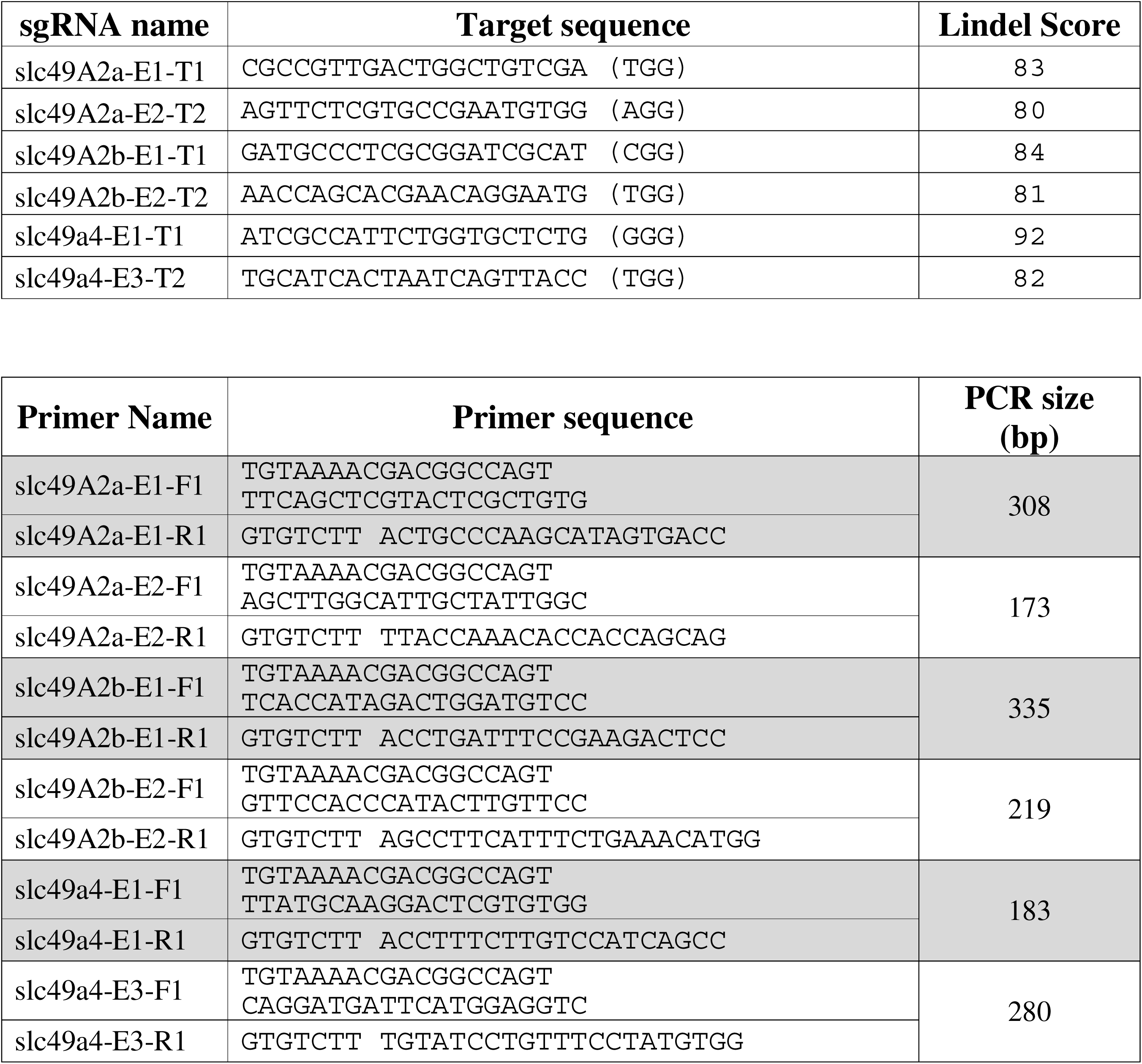
Guide RNAs, primers and target sequences used for CRISPR/Cas9 in zebrafish.

