## Supplementary figures and images for "A conserved vesicular circuit enables metabolic adaptation and maintains systemic heme homeostasis"

### FigS1

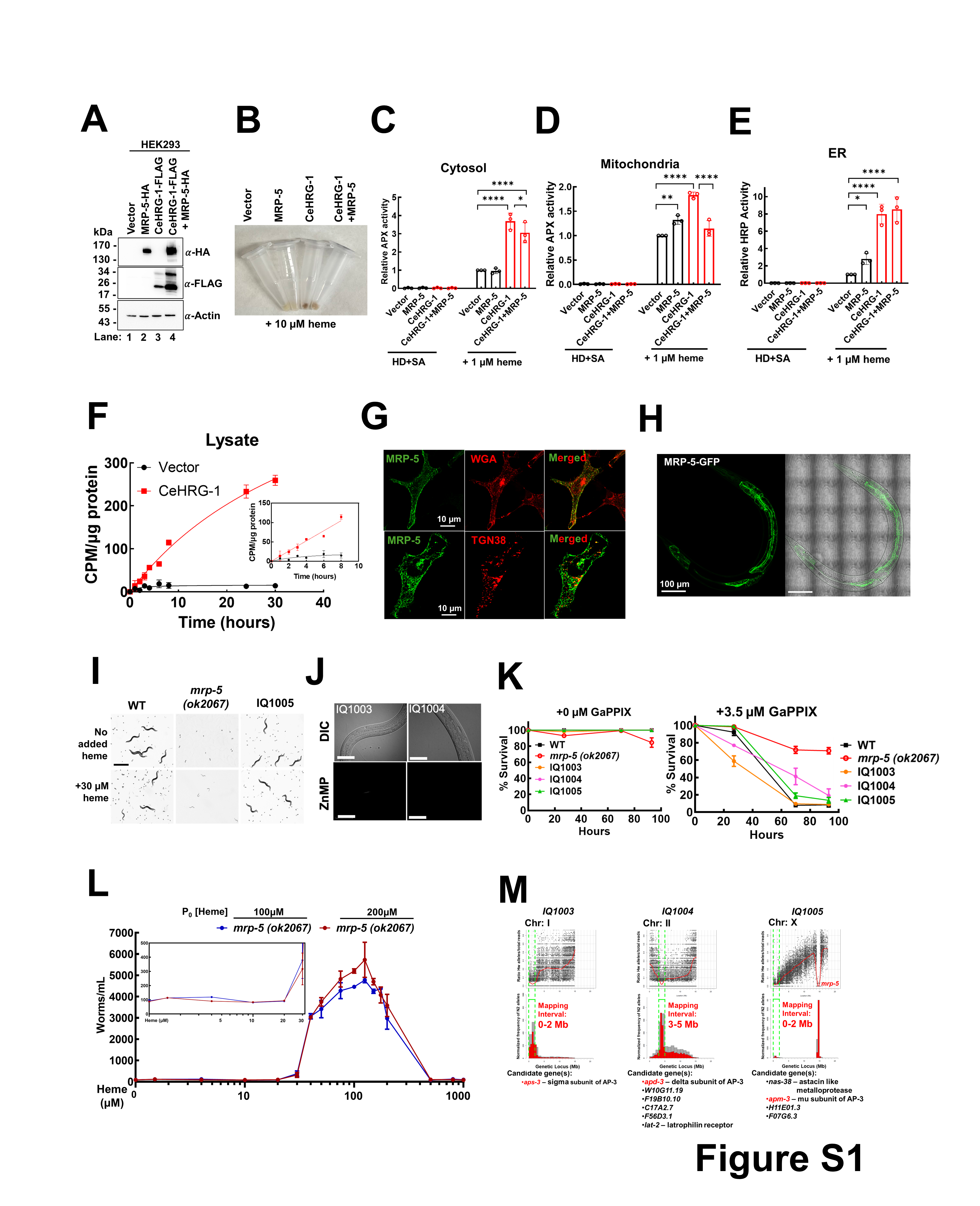

### FigS1.2

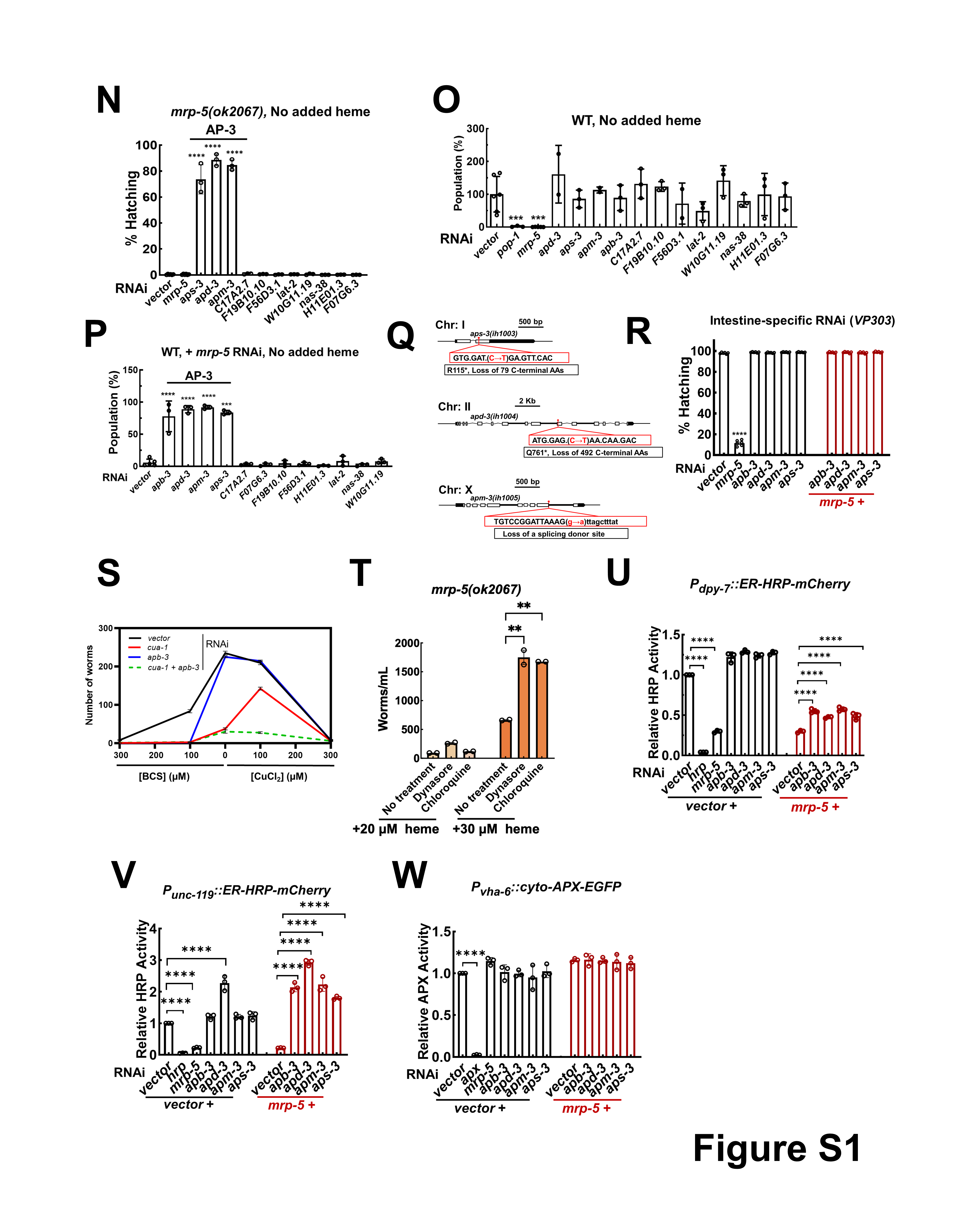

### FigS2

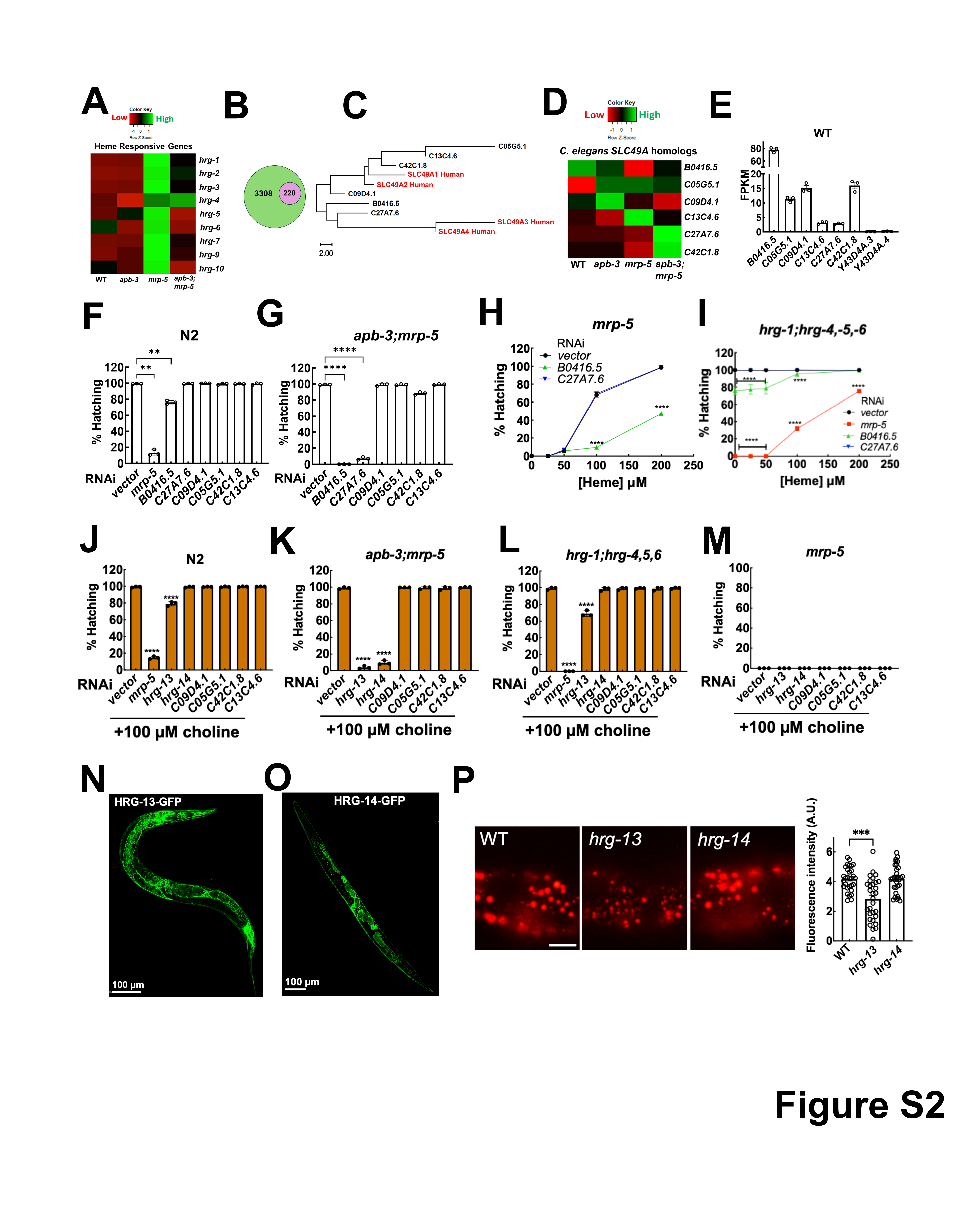

### FigS3

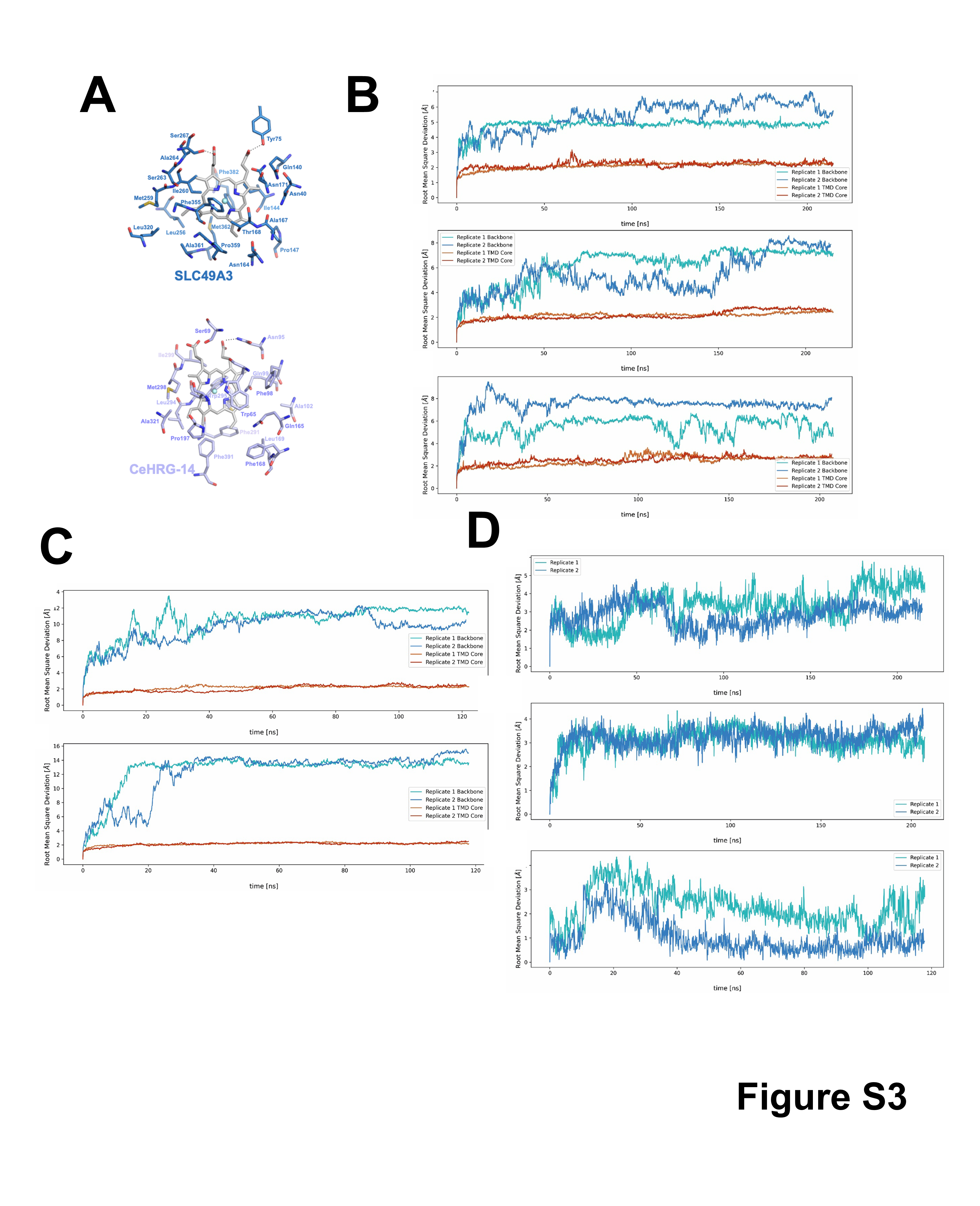

### FigS3.2

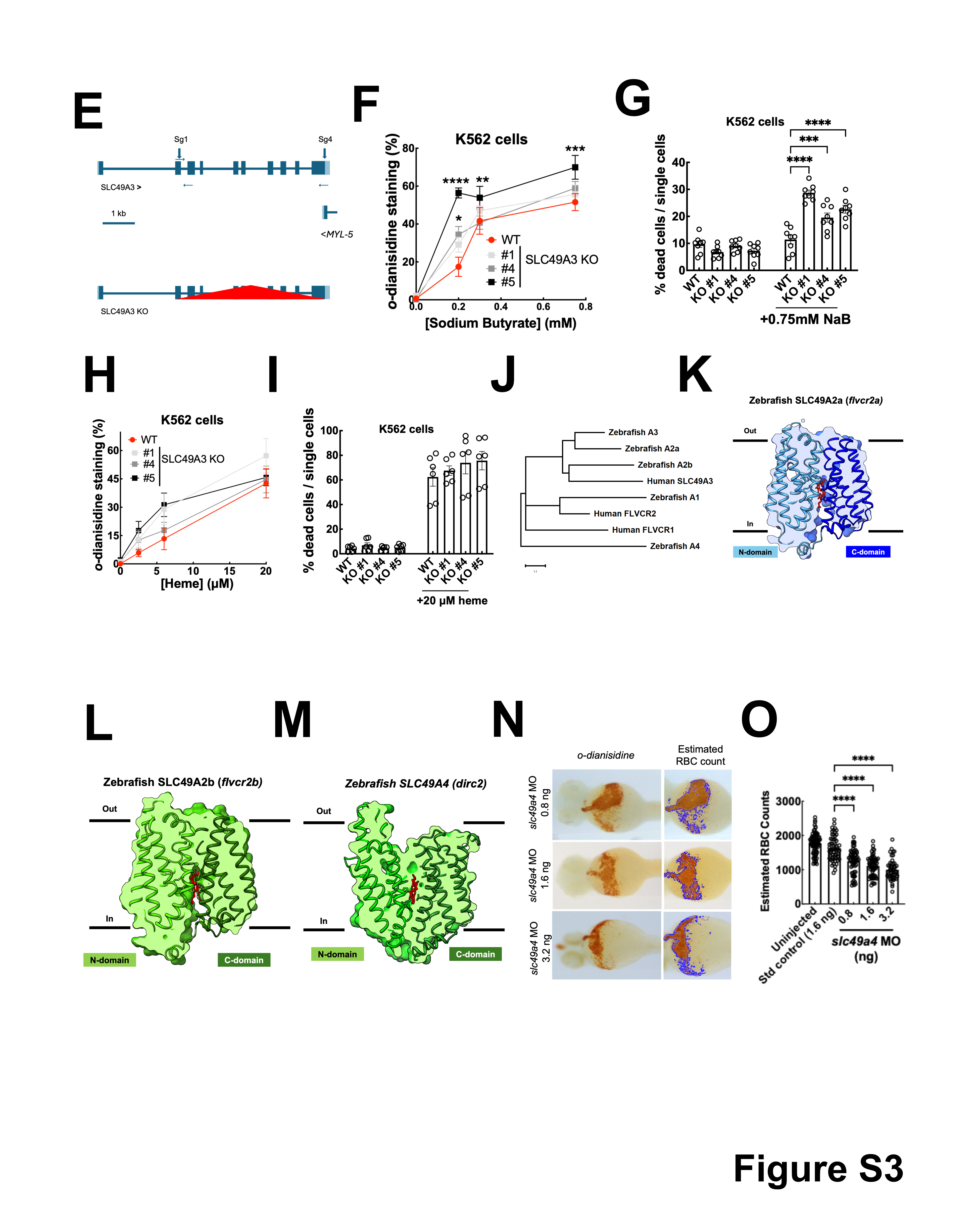

### FigS4

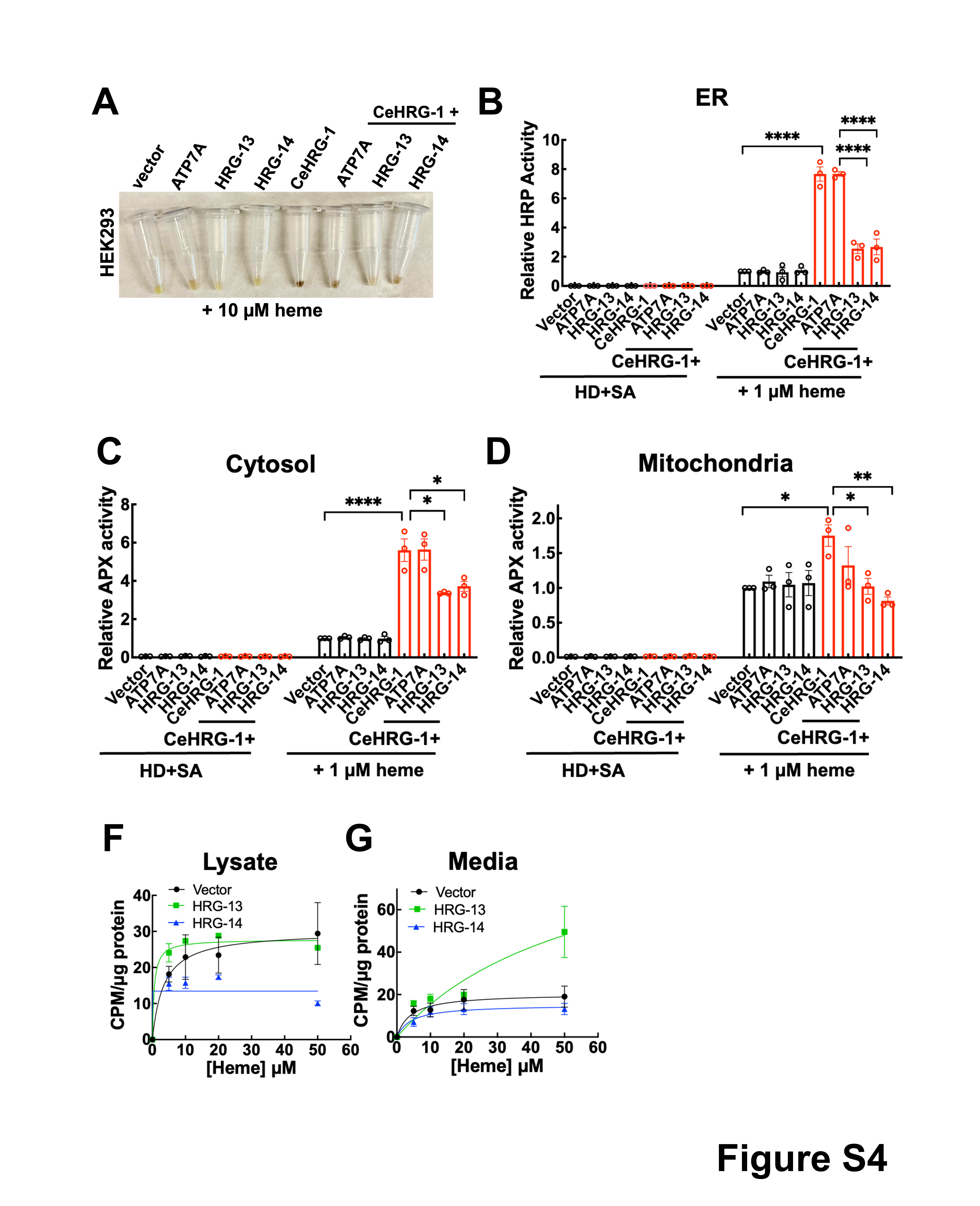
